# A tale of two transcriptomic responses in agricultural pests via host defenses and viral replication

**DOI:** 10.1101/2020.09.25.312561

**Authors:** Pramod Pantha, Subbaiah Chalivendra, Dong-Ha Oh, Bret Elderd, Maheshi Dassanayake

## Abstract

**Background:** *Autographa californica* Multiple Nucleopolyhedrovirus (AcMNPV) is a baculovirus with a high potential for its use as a biopesticide against arthropod pests. The budded form of the virus causes a systemic infection when it escapes the midgut to enter the hemolymph of susceptible hosts. Yet, the specific molecular processes underlying the biocidal activity of AcMNPV on its insect hosts are largely unknown.

**Results:** In this study, we describe the transcriptional responses in two major pests, *Spodoptera frugiperda* and *Trichoplusia ni*, to determine the host-pathogen responses during AcMNPV infection, concurrently with the viral response to the host. We assembled species-specific *de novo* reference transcriptomes of the hemolymph to identify key transcripts that respond during pathogenesis in these arthropod models where genomic resources are sparse. We found that the suppression of transcriptional processes related to chitin, a metabolite critical for basement membrane stability and tracheal development are central in establishing a systemic infection. Synergistic transcriptional support was observed to suggest suppression of immune responses and induction of oxidative stress indicating disease progression in the host. The entire AcMNPV core genome was expressed in the host hemolymph and viral genes predominantly associated with the budded virus replication, structure, and movement were more abundant than those associated with the occlusion-derived virus. Genes known to directly arrest host cell cycle and development were among the most abundant AcMNPV transcripts in infected hosts. Interestingly, several of the host genes (e.g. *Chitin synthase*) that were targeted by the pathogen as revealed by our study are also targets of several chemical insecticides currently used commercially to control arthropod pests.

**Conclusions:** Our results reveal an extensive overlap between biological processes represented by genes differently expressed in both hosts, as well as convergence on highly abundant viral genes expressed in the two hosts, providing an overview of the host-pathogen transcriptomic landscape during systemic infection. Given the diversity of AcMNPV strains that infect a wide range of insect hosts, our study provides a framework where pathogen strains could be selected to target specific host genes that facilitates modulation of the infection strength and specificity of the susceptible hosts.

## Background

Baculoviruses are ubiquitous in nature and affect a wide-range of insects [1]. These highly virulent viruses are arthropod-specific and mainly infect lepidopteran larvae [2]. Baculoviruses belong to the family Baculoviridae and have large rod-shaped nucleocapsids with circular DNA genomes [3–5]. An outer lipoprotein envelope surrounds one or more nucleocapsids to form a virion which are themselves bundled together within a protein matrix to form an occlusion-derived virus [3]. Occlusion-derived viruses are large enough to be seen and quantified using a hemocytometer under a light microscope [6]. In lepidopteran populations, baculovirus epizootics begin when a larva consumes virus-contaminated foliage [7]. If enough virus is consumed, a fatal infection occurs. The virus replicates within the larva until the virus triggers the liquefaction of the insect host, which releases occlusion-derived viruses onto nearby foliage [7]. After the virus is released, uninfected larvae eat the newly contaminated foliage and the cycle continues. Overtime, occlusion-derived viruses degrade due to exposure to ultra-violet light [8].

Baculoviruses usually have specific host ranges and most of them only infect congeneric insect species [1]. The most notable exception is the *Autographa californica* Multicapsid Nucleopolyhedrovirus (AcMNPV), which infects over 35 species belonging to 11 lepidopteran families [9]. The unusually broad host range for a baculovirus has made AcMNPV one of the most promising candidates for bioinsecticide development. AcMNPV is also widely used as a molecular tool in gene delivery systems and for engineered protein production in insect cell cultures [10–13]. Several strains of AcMNPV have been sequenced [14–16]. Their genomes are ∼134 kbp in size and contain ∼150 tightly spaced genes [14–16]. Due to the host-specific virulence of individual strains, AcMNPV is a potent biopesticide in integrated pest management systems that could spare beneficial insects specially in ecologically sensitive areas[17, 18].

Baculovirus results in two distinct virion phenotypes upon infection in insect hosts [5, 19]. First, the occlusion-derived virus is transmitted among insects primarily via horizontal transmission when uninfected hosts inadvertently consume the virus. This will often result in a lethal infection [6]. Second, following infection of the midgut epithelial cells, the budded virus causes secondary infection in the open circulatory system and, subsequently, invades cells in other tissue types [20]. Besides horizontal transmission, vertical transmission between mother and offspring may also occur. However, vertical transmission often results in a “covert” infection that does not kill the host [7].

The fall armyworm (*Spodoptera frugiperda*) and the cabbage looper (*Trichoplusia ni*) are among major agricultural pests vulnerable to AcMNPV infection. These two pests together pose a significant threat to global food security, affecting over 150 crops including corn, sorghum, rice, sugarcane, soybean, and cotton [21, 22]. The total yield loss by *S. frugiperda* alone in 12 maize producing African countries in 2017 was estimated to be between US$2.48 and $6.19 billion [22]. If appropriate control measures are not applied, these pests together can exacerbate the problem of food security and livelihood of many small farmers worldwide due to their wide host range. They are difficult to control due to their rapid spread and the development of their resistance to many insecticides [23–25]. Therefore, AcMNPV strains that can naturally infect these serious agricultural pests offer a promising mode of pest control. However, it is imperative to understand the mode of infection, disease progression, and epidemiology of a naturally occurring virus before its commercialization, to minimize unintentional secondary effects [25].

Both host species are widespread multivoltine (i.e., multiple generations per year) pests that attack a number of crops throughout North and South America [26–28]. Females lay eggs in large clusters consisting of hundreds of individuals [27, 29]. After hatching, *S. frugiperda* has six larval instars or development stages before the larvae pupate and later emerge as adults; whereas, *T. ni* has five larval instars [30, 31]. These two pests are readily infected in nature by baculoviruses, particularly when they reach large population densities [2, 32]. A typical disease outbreak or epizootic occurs when recently hatched first instars or neonates consume contaminated leaf tissue or egg casings [6]. Once an individual larva is infected, the larva does not continue to grow or molt to larger instars; whereas, uninfected individuals do.

*In vivo* studies investigating the genetic basis for AcMNPV infection and the integrated host responses are quite limited. Most studies exploring transcriptional regulation of these host-pathogen interactions use cell cultures infected with the virus. The transcriptome responses of *S. frugiperda* [33, 34] and *T. ni* [4, 35] cell cultures infected with AcMNPV have shown quite divergent transcriptional profiles, which makes it difficult to deduce the impact of these responses in intact organisms. Recently, Shrestha et al., (2019) described the *in vivo* transcriptional response of *T. ni* during AcMNPV infection. They reported the oral to midgut tissue-specific transcriptomic responses at the primary stage of infection in 5^th^ instar larvae. *In vivo* studies that explore the transcriptional dynamics in response to AcMNPV infections appear to be even fewer in *S. frugiperda*. To our knowledge, studies exploring the gene expression profile of the AcMNPV during its infection of intact hosts along with the dynamics in host transcriptomes are also absent either in *S. frugiperda* or *T. ni*.

In this study, we report the host-pathogen transcriptional responses of the early systemic infection phase. The transcriptional profiles in the host hemolymph capture host responses to the virus as well as the viral responses to the hosts. Our results indicate major transcriptional changes to support initiation of critical cellular and developmental adjustments in the host during pathogenesis.

## Results

### Lethal effects of AcMNPV on *S. frugiperda* and *T. ni*

Clearly, both *S. frugiperda* and *T. ni* were adversely affected by increased doses of AcMNPV (Fig. 1a and b) resulting in the death of a large proportion of larvae at higher doses. *S. frugiperda* required a much larger dose of the virus to become infected as compared to *T. ni* (Fig. 1a and b). This was further demonstrated by the fact that the median LD95 for *S. frugiperda* was over a magnitude higher than the LD95 for *T. ni* (Fig. 1c). Given the relatively good fit of the logistic model to the data and the relatively narrow credible intervals, the median LD95 for both species was reasonably well estimated.

**Figure 1.**
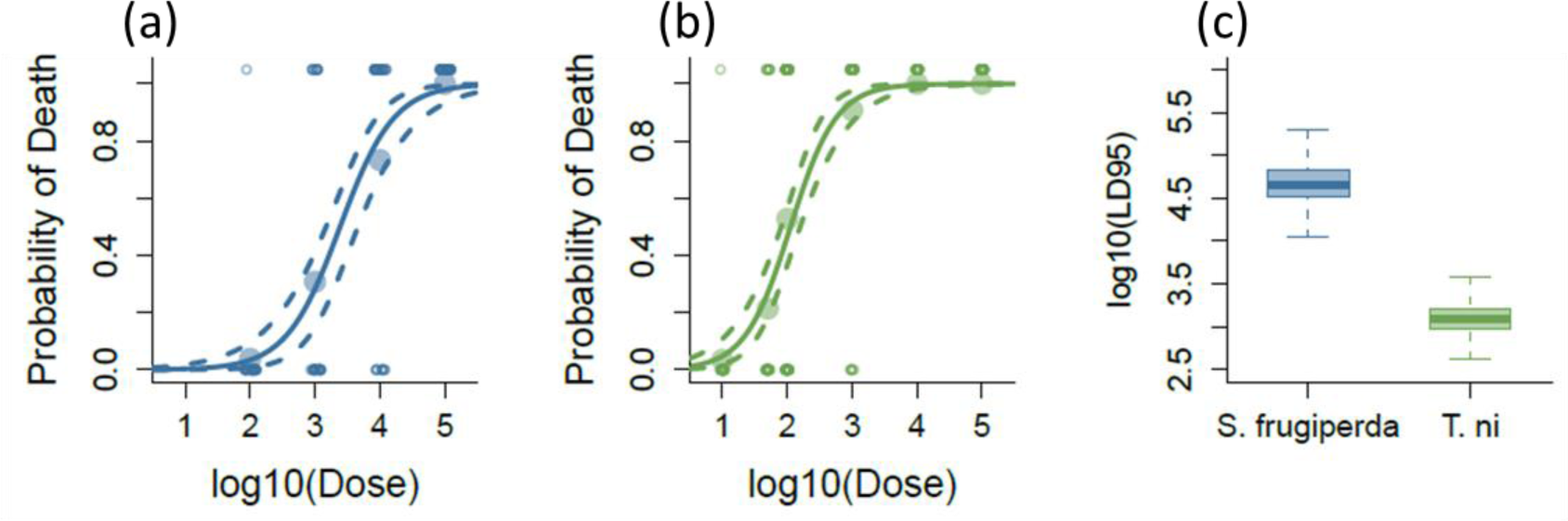
Lethal AcMNPV dose determination for *Spodoptera frugiperda* and *Trichoplusia ni.* The effects of increasing doses of baculoviruses on probability of larval death for [a] *S. frugiperda* and [b] *T. ni* along with the corresponding [c] box plot of the lethal dose at which 95% of the individuals would be expected to die identified as LD95. For [a] and [b], the solid line is the median dose-response curve and the dashed lines are the 95% credible intervals for the curve. The large filled points represent the mean response for each dose and the small open points are the individual data. These data are jittered for ease of presentation. For [c] the dark line of the box plot is the median with the box encompassing the interquartile range between the first and third quartiles and the whiskers represent 1.5 times the interquartile range.

### *De novo* assembly and annotation of *S. frugiperda* and *T. ni* 4^th^ instar reference transcriptomes

We report the most curated reference transcriptomes that represent the hemolymph tissue of *S. frugiperda* and *T. ni* currently available. On average, 62 million raw reads were obtained for each RNA-seq sample generated for *S. frugiperda* and *T. ni* (Supplementary Table 1). The fully assembled transcriptomes are available at NCBI BioProject PRJNA664633.

We selected 17,908 *S. frugiperda* transcripts (mean length 1,458 nt) and 19,472 *T. ni* transcripts (mean length 1,773 nt) to represent the protein-coding reference transcriptomes (Table 1). The number and length distribution of total protein-coding transcript models in the current reference transcriptomes (Supplementary Fig. 1a and b) were comparable to the protein-coding transcripts available for *Bombyx mori* [37], *Helicoverpa armigera* [38], *Spodoptera litura* [39] and the genome of *T. ni* [40] (Supplementary Fig. 1c and d). Our predicted protein coding transcripts mainly contained complete ORFs with start and end codons included in the transcript model (Supplementary Fig. 2a). We were able to map >75% of the initial RNA-seq reads to the reference transcriptomes for both species (Supplementary Table 2).

**Table 1.** Summary of *de novo* assembled reference transcriptomes of *S. frugiperda* and *T. ni*.

| Transcriptome assembly features | *S. frugiperda* | *T. ni* |
| --- | --- | --- |
|  | Coding <sup>a</sup> (non-coding <sup>b</sup>) | Coding <sup>a</sup> (non-coding <sup>b</sup>) |
| Total assembled transcripts | 17,908 (101,169) | 19,472 (147,934) |
| Percent GC | 42.68 (35.28) | 41.16 (35.24) |
| Contig N50 (nt) | 2,279 (532) | 2,955 (656) |
| Average contig length (nt) | 1,458 (495) | 1,773 (549) |
| Smallest contig length (nt) | 297 (224) | 297 (201) |
| Longest contig length (nt) | 29,765 (14,343) | 30,823 (23,482) |
| Number of ORFs | 28,433 (-) | 31,292 (-) |
| Average ORF length (nt) | 920.28 (-) | 841.86 (-) |
| Smallest ORF length (nt) | 297 (-) | 297 (-) |
| Longest ORF length (nt) | 27,558 (-) | 27,408 (-) |
| Mapped sequenced read % to the reference assembly | 76 | 84 |
| Detected complete BUSCOs (%)<sup>c</sup> (Arthropoda reference) | 80.6 | 82.1 |

<sup>a</sup> represents transcript models with a predicted open reading frame (ORF)
<sup>b</sup> represents transcript models without a predicted ORF
<sup>c</sup> BUSCO (Benchmarking Universal Single-Copy Orthologs) v1.22 (Simão et al., 2015)

As *D. melanogaster* genes provided the most amount of functional attributes available for an arthropod model, we first annotated 5,878 *S. frugiperda* and 6,219 *T. ni* transcript models based on the *D. melanogaster* reference models where possible (see methods). The NCBI insect-Refseq database was used to annotate another 9,273 transcripts from *S. frugiperda* and 9,751 transcripts from *T. ni* (Supplementary Fig. 3a). The remaining transcripts were subjected to BLATX against the NCBI-nr databases to annotate 1,278 *S. frugiperda* and 888 *T. ni* transcripts. A final pool of remaining transcripts that did not show convincing similarity to other known eukaryotic transcripts (1,479 *S. frugiperda* and 2,614 *T. ni* transcripts) were annotated as “unknown putative proteins”.

We assessed the completeness of the reference transcriptomes based on the expected presence of core genes in metazoans as identified by the BUSCO database [41]. *S. frugiperda* and *T. ni* reference transcriptomes were found to have 87.3% and 87.2% expected BUSCOS respectively, suggesting that these transcriptomes contain a core gene component comparable to the high quality lepidopteran genome model of silkworm [42] (Supplementary Fig. 2b). Furthermore, our *S. frugiperda* reference transcriptome showed a better BUSCO representation than the previously published *S. frugiperda* genome and transcriptome assemblies [43, 44] (Supplementary Fig. 2b). Only 36% of RNA-seq reads generated for *T. ni* in our study mapped to a genome assembly recently made available for this insect [40], compared to the 84% of mapped reads to our reference transcriptome. These comparisons confirm the appropriateness of the use of our reference transcriptomes for our downstream analyses.

### Host transcriptomic responses to the AcMNPV infection

We identified 175 *S. frugiperda* differently expressed transcripts (DETs) and 138 *T. ni* DETs in response to the AcMNPV infection (Fig. 2a and b). The DETs represent ∼1% *S. frugiperda* and ∼0.7% *T. ni* of respective reference transcriptomes. The relatively small sets of differently co-expressed genes suggest that the observed transcriptomic response pertains to an active host responding to the infection, rather than largely missregulated transcriptomes represented in a dead or a dying host overrun by the pathogen. In addition, the transcriptional responses between control and infected larvae within a host species differed minimally relative to the differences between basal transcriptomes of the two hosts (Supplementary Fig. 4). The wide divergence observed in the basal transcriptomes of the two host species is not surprising, since they belong to two different genera.

**Figure 2.**
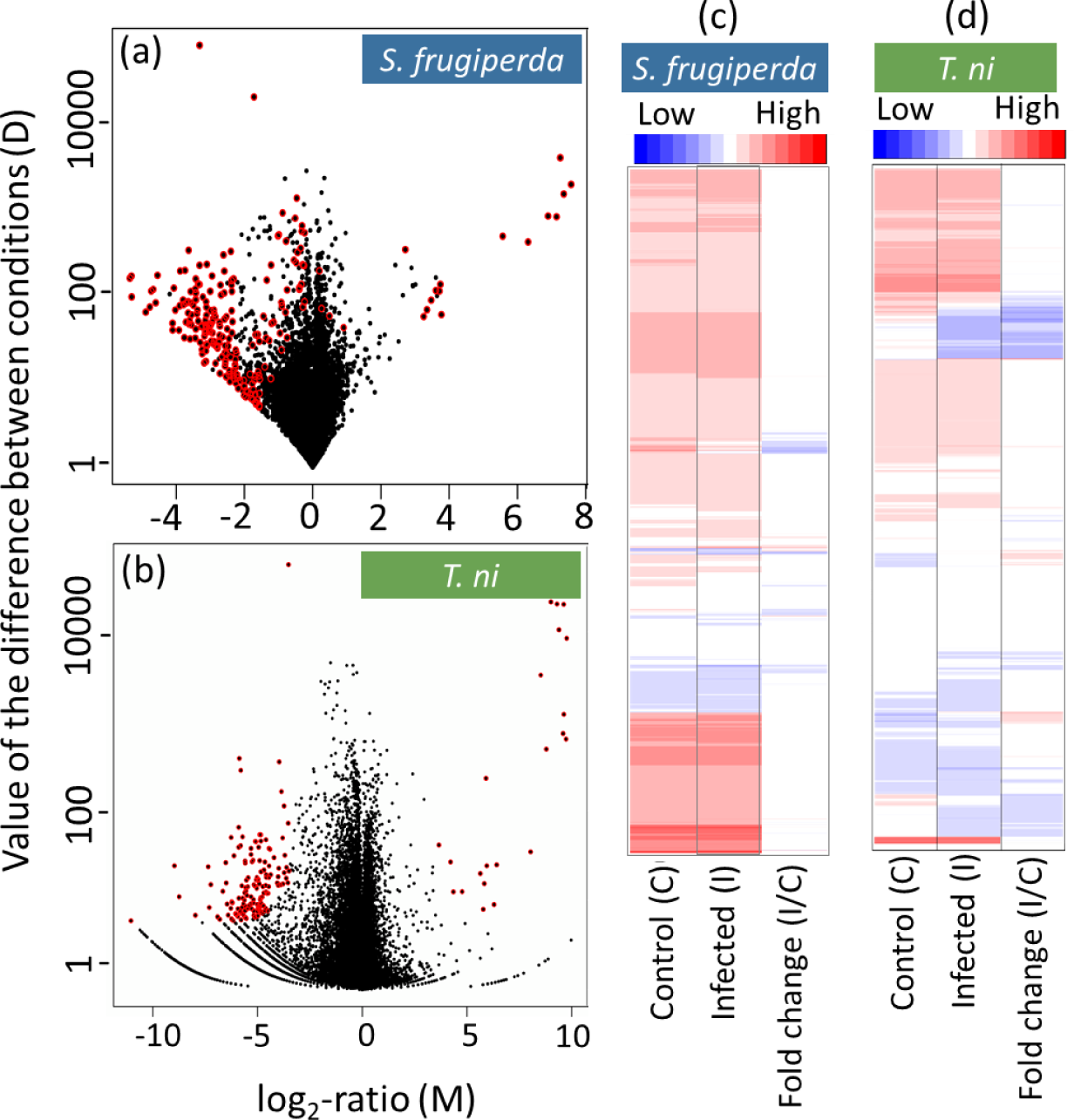
Host transcriptomic response to AcMNPV infection. Summary MD plots of the normalized expression values for control and AcMNPV treated samples for [a] *S. frugiperda* coding transcripts (18 up- and 157 down-regulated transcripts) and [b] *T. ni* coding transcripts (20 up- and 118 down-regulated transcripts). Differently expressed transcripts (DETs) at a q-value cutoff of 0.95 are indicated in red dots. All the DETs with their respective fold changes are listed in the Supplementary Table 4. Heatmaps show log2 normalized expression of control, AcMNPV-infected, and log2 fold changes of 17,873 transcripts for *S. frugiperda* [c] and 18,203 transcripts of *T. ni* [d]. The genes are clustered based on their expression strength similarity.

Our results show that, in both host species, transcripts suppressed due to infection differed by orders of magnitude compared to those transcripts that were induced by viral infection (Fig. 2). This is consistent with the trend observed in previous transcriptomic studies in cell cultures of AcMNPV-infected *S. frugiperda* and *T. ni* [33, 34, 36, 45, 46]. Interestingly, we see extensive similarities across multiple biological processes as deduced from the functional attributes of DETs in each species, suggesting a shared host response to the AcMNPV pathogen. Overall, 83.4% *S. frugiperda* and 89.1% *T. ni* DETs could be assigned to functionally informative annotations. This was based on either functional validation of a putative homolog in *D. melanogaster* or a homolog reported with a putative function in another lepidopteran host. The number of Gene Ontology (GO) annotations were used when available but was more limited as GO annotations largely depended on the sequence similarity of *S. frugiperda* and *T. ni* transcript models to a *D. melanogaster* gene that also had an assigned GO term. In the following sections, we highlight the shared host transcriptomic responses via enriched functional processes based on clustering of functional annotations of DETs. All DETs with functional annotations that had a fold change of 4 or more in response to the AcMNPV infection were considered. The full list of DETs and their assigned GO terms (when available) are presented in Supplementary Table 5.

### Chitin metabolism and epithelial membrane associated processes were suppressed in AcMNPV-infected hosts

The two largest enriched functional clusters out of six in *S. frugiperda* and the largest cluster of the two in *T. ni*, represented in the “suppressed” set reveal a coordinated downregulation of chitin-related genes (Fig. 3). Chitin metabolism and its associated pathways are central to the formation and stability of the extracellular matrix, basement membrane, cuticle, and the tracheal system that are in close contact with the hemolymph tissue. The genes associated with this cluster are not limited to those with assigned GO terms (Fig. 4b and Supplementary Table 4). Among these, genes associated with chitin synthesis (*chitin synthase 1/kkv*); genes encoding chitin-binding proteins specially in the peritrophic matrix (*Gasp*) [47, 48]; other genes known for their chitin associated functional roles in cuticle development such as the *Osiris* gene family members and *Dusky-like* (*Dyl*) that regulate the deposition of chitin on bristles were significantly down-regulated in *S. frugiperda* (Supplementary Table 4) [49–52]. Interrupted chitin metabolism at the cellular level is tightly coupled to the organ integrity, particularly of the midgut and the tracheal system. Drosophila *chs1* mutants with suppressed expression also show defective tubular structure, irregular tracheal epithelial tube expansion, and irregular subapical cytoskeletal organization [53]. The host genes *serpentine* (*serp*) and *vermiform* (*verm*) that bind to chitin and modify its surface play significant roles in the tracheal tube development [54–56] together with *uninflatable* (*uif*) that regulates tracheal growth and molting [57]. These and many other cuticle and tracheal growth related genes were highly co-suppressed in infected host tissue (Fig. 4b and Supplementary Table 4).

**Figure 3.**
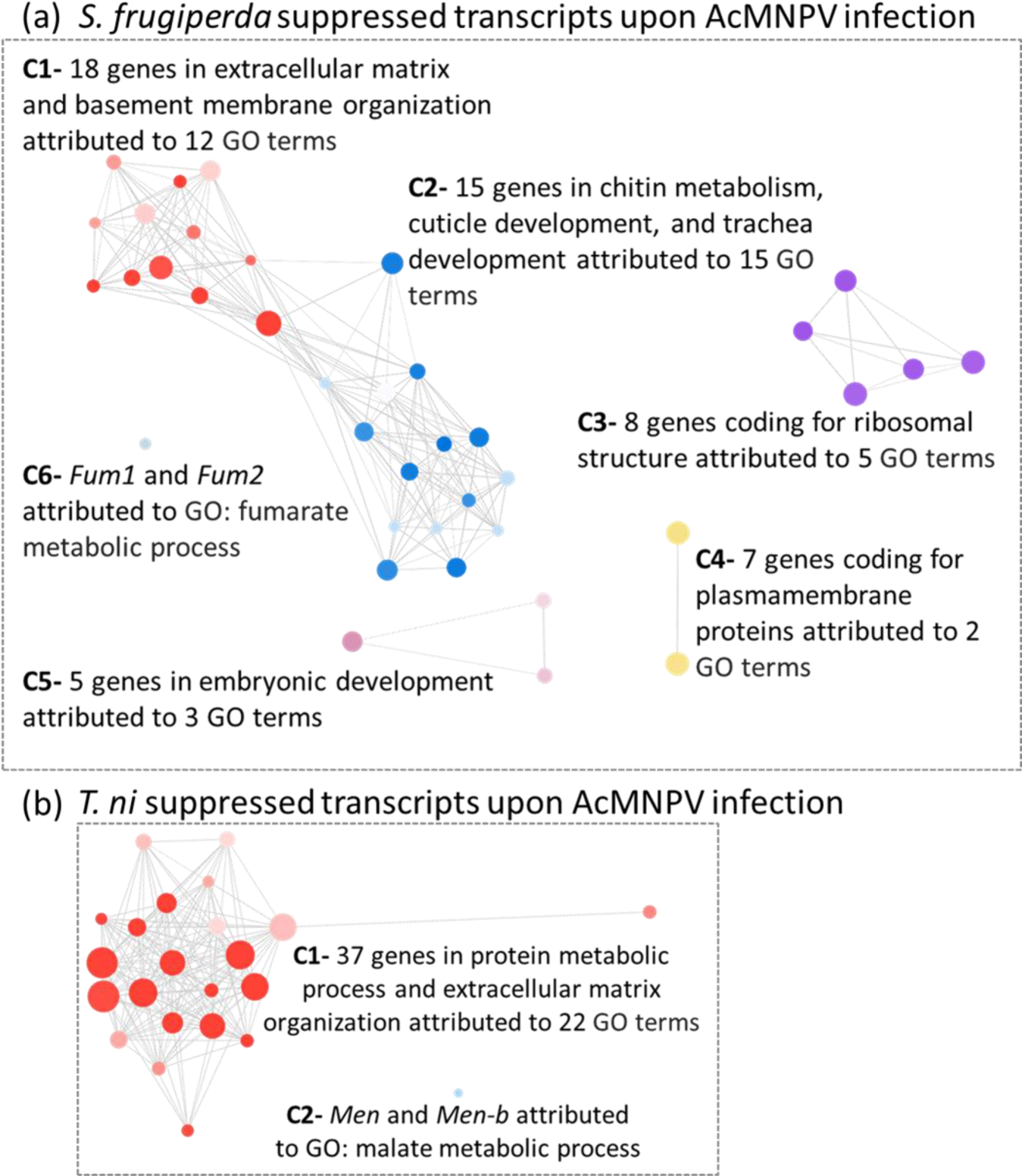
Overview of enriched functional processes represented by suppressed genes in the infected host transcriptomes. Functional clusters of *S. frugiperda* [a] and *T. ni* [b] transcripts suppressed upon AcMNPV infection. Six and two distinct functional clusters were identified for *S. frugiperda* and *T. ni* respectively. The network connect GO terms, marked as nodes connected by edges that represent a minimum overlap of 80% genes (in the smaller GO term of the pair) based on Markov clustering (MCL). Distinct colors indicate shared functional groups within the network. The radius of the node represents the number of genes and the shade represents FDR adjusted p-value of ≤0.05 enrichment assigned using GOMCL [216]. Each cluster is named based on the largest enriched GO term in a given cluster.

**Figure 4.**
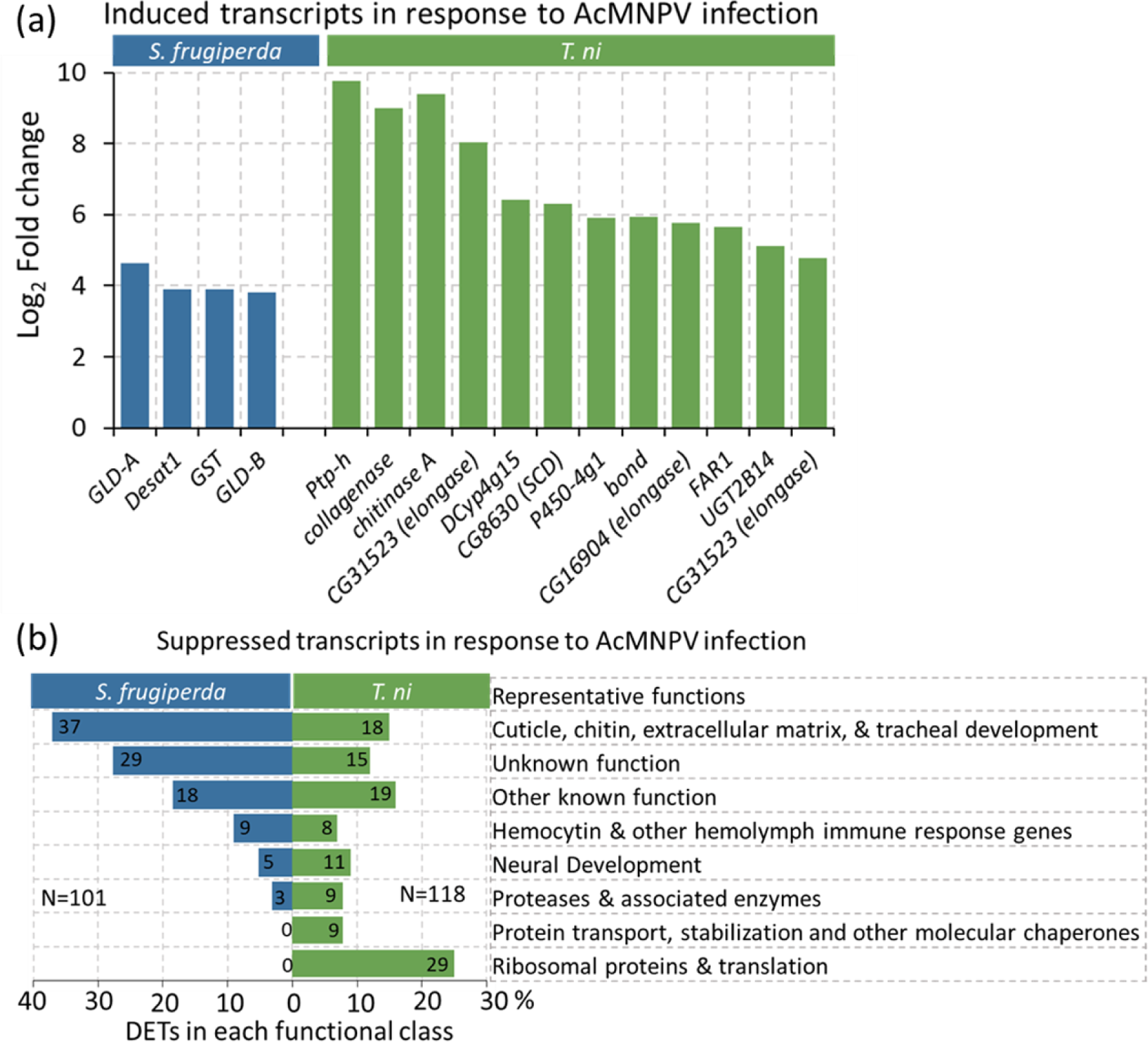
*S. frugiperda* and *T. ni* differently expressed transcripts (DETs) in response to the AcMNPV infection. Induced DETs are shown in [a] and summarized groups that represent a total of 101 in *S. frugiperda* and 118 suppressed DETs in *T. ni* are given in [b].

The budded virus exiting the midgut epithelial cells needs to penetrate the basement membrane of the gut epithelium before entering the hemocoel and then the basement membrane of tracheal cells for systemic infections [56]. Collagen is a fundamental component of basement membranes of both gut and tracheal epithelia [58, 59]. Therefore, genes associated with collagen metabolism and other integral components of the basement assembly are expected candidates for virus regulated transcriptional processes in the host. Transcripts coding for structural components of the extracellular matrix including collagen were among the most significantly suppressed in response to the AcMNPV infection in both species (Fig. 4b and Supplementary Table 4).

We observed multiple transcripts associated with glycoproteins, likely formed in hemocytes that function in basement membrane stability, highly suppressed coordinately in both species during AcMNPV infection. Among them, *laminins*, *osteonectins* (*SPARC*), and *papilins* are notable. Laminin is the most prevalent glycoprotein in the basement membrane and is also found in extracellular matrices of tracheal cells. It is formed of three chains coded by *LanA*, *LanB1,* and *LanB2* [60–62]. Notably, we found transcripts that represent all three *Laminin* chains to be coordinately down-regulated in the infected tissue in both species (Supplementary Table 4). *SPARC,* known as a Ca^2+^ binding extracellular glycoprotein that modulates cellular interactions with the extracellular matrix [63] was also down-regulated in both hosts during the budded virus infection stage (Supplementary Table 4). *SPARC* is particularly expressed during cellular injury or wounding that require tissue remodeling [64] and functions in basal lamina assembly and stability [65, 66]. Similarly, transcripts potentially coding for papilins were co-suppressed in infected samples of both hosts (Supplementary Table 4). Papilins expressed in hemocytes are a prominent group of sulfated glycoproteins that contribute to basement membrane structure [67–69]. The coordinated suppression of chitin and basement membrane associated glycoproteins in our results indicate a strong transcriptomic signal for weakened membrane stability in infected host tissue during the budded virus invasion into the hemocoel.

### Transcripts associated with hemocyte-induced defenses and immune responses were suppressed during systemic infection

Membrane damage in contact with the hemocoel is sensed by hemocytes and these can initiate immune responses during pathogen invasions. Melanization is a major hemocyte-driven defense response that leads to blood clotting. Surprisingly, this pathway appeared to be suppressed as evident from the down-regulation of multiple host genes in both species in response to the AcMNPV infection. *Hemocytin* is a key gene that mediates hemocyte aggregation and hemolymph melanization in lepidopteran innate immunity against pathogens [70–72]. *Hemolectin* is specifically expressed in larval hemocytes, and acts as a clotting factor involved in hemostatis [73–75]. It is also known to initiate immunity responses during pathogen infections [74, 76] and is thought to play a vital role in encapsulating foreign substances during metamorphosis in *B. mori* [75]. Hemocytins and hemolectins were among the most highly suppressed genes in both *S. frugiperda* and *T. ni* infected samples (Fig. 4b and Supplementary Table 4).

Hemolymph proteases are known for their pivotal roles in defense responses against many pathogens as well as in development processes such as molting [77, 78]. The specific regulatory pathways of many of these proteins are not definitive yet, but their collective role as a functional group in insect immunity and development are established. We found multiple proteases in both infected hosts highly suppressed as a prominent group among all suppressed transcripts (Fig. 4b and Supplementary Table 4).

### Lipid metabolism and oxidative stress emerge as the most prominent functional processes induced by both hosts in response to infection

The lipid biosynthesis pathways not only affect lipid membranes, but also many other primary biological processes related to energy metabolism and signaling pathways. Interestingly, *Desaturase1 (Desat1*) is induced in *S. frugiperda* upon AcMNPV infection (Fig. 4a). Desat1 is reported to be tightly regulated at the transcriptional level [79] and is required for the biosynthesis of unsaturated fatty acids [80, 81]. Additionally, several fatty acid modification enzymes, e.g. elongases like *jamesbond/bond*, and *CYP4G*, a cytochrome P450 that performs oxidative decarbonylation of long chain fatty aldehydes [82–84] were co-induced in *T. ni*. It is notable that *bond* and *CG16904* together were assigned to 60 GO-terms, exemplifying their influence in multiple biological functions linked to their primary molecular functions in lipid metabolism [82–84] (Supplementary Fig. 4 and Supplementary Table 5).

Reactive oxygen species (ROS) generation and induction of oxidative stress are inevitable when host membranes are disrupted and lipid metabolism is altered during host-pathogen interactions. Supportive of this expectation, all three genes induced in the infected *S. frugiperda* hemolymph in addition to *Desat1* (i.e. above a 4-fold expression change) relate to oxidative stress (Fig. 4a). These include transcripts coding for a cytosolic *GST* and two *FAD-glucose dehydrogenases (GLD).* GSTs form a broad family of critical defense proteins against oxidative stress [85, 86] and FAD-glucose dehydrogenase can induce ROS generation as a defense response [87]. A recent study has also reported that FAD-glucose dehydrogenase is induced as a defense response during AcMNPV infections in *Helicoverpa zea* [88].

### AcMNPV genome response to the insect hosts

To check whether viral sequences were present in our hemolymph samples, we mapped RNA-seq reads from both species to the published AcMNPV genome [16] (Supplementary Table 2b). As expected, viral sequences were detected almost exclusively in the infected samples. We mapped 1.13% and 7.41% of total reads from infected *S. frugiperda* and *T. ni* samples, respectively, to the AcMNPV genome. It was interesting that a small number of reads from *T. ni* control samples (<0.01%) were mapped to the AcMNPV genome (Supplementary Table 2b). While it is not conclusive that these could represent domesticated viral genes expressed at low levels in the *T. ni* genome, previous studies have indicated that AcMNPV genes are found in arthropod genomes as a result of horizontal gene transfer [89, 90].

The AcMNPV strain E2 genome has 149 protein-coding genes [16]. We detected 148 genes in our viral transcriptome expressed in the hemolymph (Fig 5a and Supplementary Table 6). These transcripts were categorized into nucleocapsid-associated and envelope-associated genes. Each of these two categories was further divided into their contribution to the formation of the occlusion-derived virus, budded virus, or their involvement in the formation of both virion types, following Blissard and Theilmann (2018). Viral genes related to the formation of the budded virus showed higher expression than those involved in the production of occlusion-derived virus in both nucleocapsid- and envelope-associated categories in samples from both hosts (Supplementary Fig. 6a & b and Supplementary Table 6). Budded virus compared to the occlusion-derived virus is the dominant form expected in the hemocoel during the systemic infection phase [5]. All viral genes showed higher levels of expression in infected hemolymph of *T. ni* compared to that of *S. frugiperda* (Fig. 5a and Supplementary Table 6).

**Figure 5.**
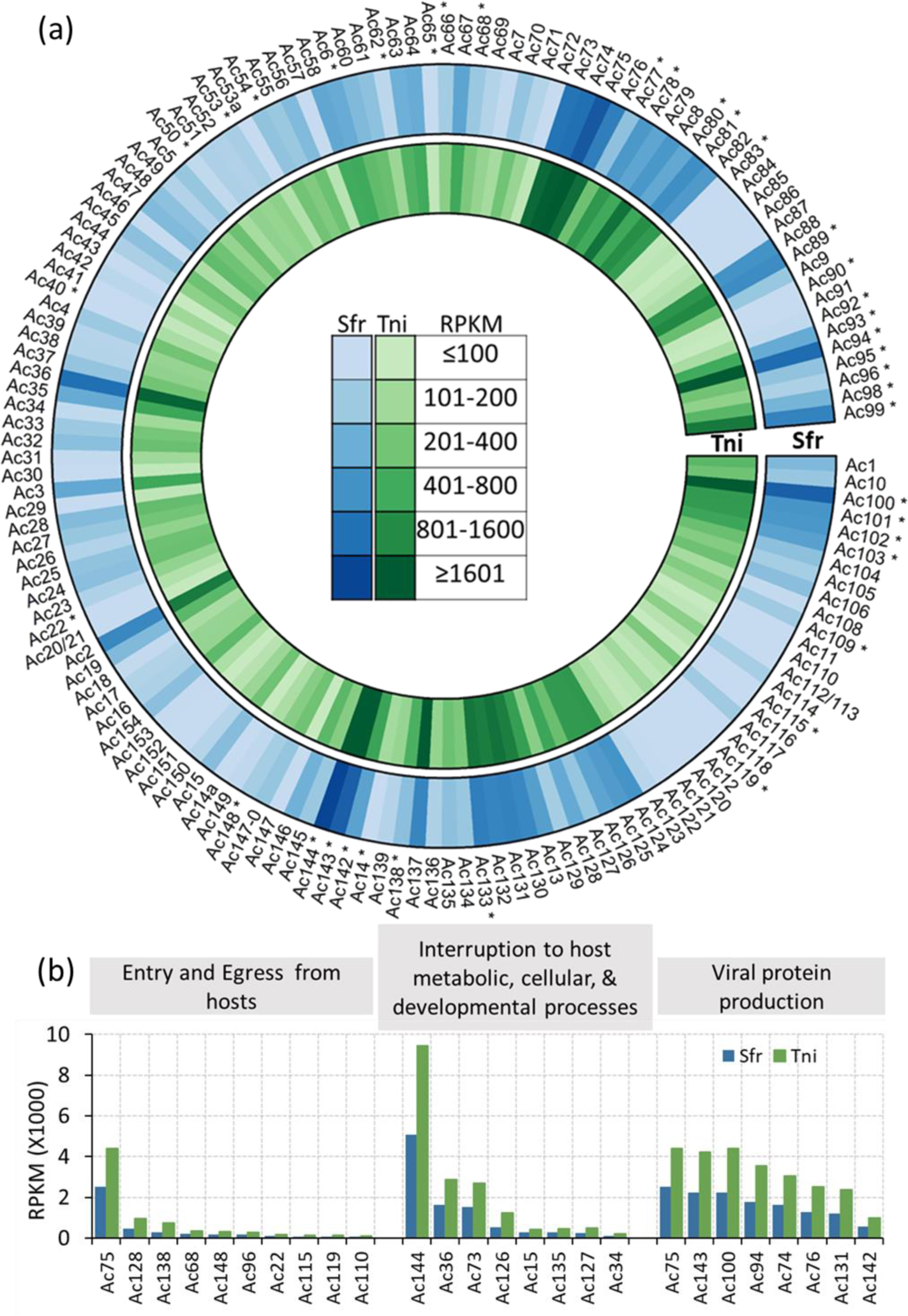
The AcMNPV genome expressed in the host hemolymph. [a] The circular plot show the normalized gene expression of AcMNPV genes in infected *S. frugiperda* and *T. ni*. Core baculovirus genes are marked with asterisks. [b] Expression of AcMNPV genes associated with entry and egress from insect hosts, interruption to host metabolic, cellular, and developmental processes, and viral protein production.

Baculovirus genes show three sequential stages of expression, marked as early, late, and very late. The early viral genes are transcribed by RNA polymerase II of the host. A unique feature of baculoviruses compared to other nuclear-replicating DNA viruses is that these viral genomes encode a DNA-directed RNA polymerase. This RNA polymerase transcribes the late and very late viral genes [91]. In the infected hemolymph tissue, we found viral genes that mark both early and late stages in their expression sequence. For example, a chromatin-like structure called the virogenic stroma is formed in the center of the nucleus of infected cells. *Ac36/pp31* is an early viral gene reported to be among the two primary viral genes that initiates this morphological change in the host cells [92, 93]. In contrast *Ac74/Bm60,* required for the budded virus production and also found in nucleocapsids of both budded and occlusion-derived virions, is thought to be expressed at a late stage [94]. Both *Ac36* and *Ac74* are among the top 10 highly expressed viral genes in infected samples of both hosts (Supplementary Fig. 6c).

Rohrmann (2013) had identified 37 core baculovirus genes that were also highly conserved in the AcMNPV genome. Half of the top 30 highly expressed AcMNPV genes in treated samples of both insect species were core genes (Fig. 5a and Supplementary Table 6). The majority of the viral transcripts in our study were associated with the production of nucleocapsid and envelope proteins. Many such integral proteins of the nucleocapsid or envelope are known to function in viral entry and exit pathways. For example, the highly expressed viral genes, *Ac75, Ac76,* and *Ac143* (Fig. 5a and Supplementary Fig. 6c) perform multiple roles associated with the formation of intranuclear microvesicles and production of the budded virus, while contributing to the structure of the occlusion derived virus envelope [95–101]. Other highly abundant viral genes that form integral components in the nucleocapsid or the envelope present in both hosts include *Ac131/ Pp34* [15, 102], *Ac142/p49* [103, 104]; *Ac94/odv-e25* [97, 98]; and Ac100/p6.9, [92, 99, 101] (Fig. 5 and Supplementary Fig. 5b). The cellular entry of the budded virus is dependent on GP64 coded by *Ac128* while the entry of the occlusion-derived virus is mediated by the family of *PIF* genes [5]. *Ac128* and the eight PIF genes (*Pif-0/Ac138, Pif-1/Ac119, Pif-2/Ac22, Pif-3/Ac115, Pif-4/Ac96, Pif-5/Ac148, Pif-6/Ac68, Pif-7/Ac110*) out of the nine members were among highly expressed viral transcripts detected in the infected host tissue in our study (Fig. 5a and b).

The viral genes affect cellular, metabolic, and developmental alterations in the host in addition to initiating viral replication and virion movement in the host cells. Three of these virus induced host metabolic processes include host membrane degradation, cell cycle arrest, and developmental arrest that stops molting. The co-expressed viral genes *chitinase* (*Ac126*) and *cathepsin* (*Ac127*) are required for the liquefaction of hosts in the late stage of infection [105, 106]. Viral chitinases act on degrading the host chitins and cathepsins are broad-spectrum proteases that degrade host tissue [107]. Both *Ac126* (found at RPKM of 540 in *S. frugiperda* and 1227 in *T. ni*) and *Ac127* (found at RPKM of 236 in *S. frugiperda* and 494 in *T. ni*) were highly exprssed in the infected samples in our study (Figure 5b and Supplementary Table 6). This indicates a strong transcriptional signal about the extensive tissue damage initiated in the host by the budded virus along with the reciprocal transcriptomic signals in the hosts that suggest interrupted membrane stability early on during the budded virus infection.

We detected compelling transcriptomic signals that suggest virus induced host cell cycle interruption, parallel to signals of host tissue detioration. *Ac144/Ac-odv-ec27* is the most highly expressed AcMNPV gene (expressed at RPKM of 5081.6 in *S. frugiperda* and 9456.6 in *T. ni*) found in infected hosts in our study (Fig. 5a). *Ac144* is an essential gene known for its role in arresting the host cell cycle at the G2/M phase [19, 104].

### Viral-host co-transcriptional interactions

Several AcMNPV transcripts and their associated proteins are known to directly interact with host proteins to regulate pathogenicity. We wanted to assess whether such host-parasite transcript interactions could be elucidated from comparing viral transcripts co-expressed with host transcripts in the infected hemolymph.

We found the budded virus-associated gene *Ac73* (RPKM of 1530.9 in *S. frugiperda* and *2694.3* in *T. ni*) (Supplementary Fig. 6c, Supplementary Table 6), that is thought to regulate host Hsp70 [108, 109] among the top 5% viral genes expressed in our study. *S. frugiperda* Hsp70 has been reported to be a required gene to express AcMNPV genes and complete the infection cycle [110]. Even though the infected hemolymph transcriptomes in our study contain the transcripts potentially coding for *Hsp70* (TR12464|c0_g1_i1, DN38479_c0_g1_i1, Supplementary Table 3), it was not significantly induced during the time of sampling (Supplementary Table 4). However, multiple transcripts coding for other molecular chaperones, protein transport, and modification associated with ER were highly suppressed in infected *T. ni* hosts (Fig. 4b).

Host lipids play multifarious roles in a virus life cycle, right from the entry of the virus into host cells by endocytosis, during replication in protected membrane vesicles, and till the virions exit the cell by exocytosis. For example, host fatty acid desaturases are required for virus replication to alter the fluidity and plasticity of membranes for viral replication complexes [111]. As described earlier, host *Desat1* along with several transcripts associated with fatty acid synthesis are upregulated in the infected hosts.

Viral entry and egress pathways highly depend on cell shape, entry and exit to the nucleus, and microvesicles regulated by host actins [90, 112]. A late viral gene, *Ac34* induces nuclear actin polymerization that promotes virus replication, and nuclear export of the virus [109, 113, 114]. In our study, *Ac34* is another highly abundant viral transcript present in the hemolymph. Reciprocally, we observed a marginal induction in *S. frugiperda Act57B* (Supplementary Table 4). *Act57B* is a major myofibrillar actin gene expressed during larval stages in Drosophila [115] and encodes a major structural protein found in the hemolymph [116]. It is unclear whether viral *Ac34* directly regulates the host *Act57B.* Previous studies have reported that *Ac34* directly regulates the host actin-associated Arp2/3 protein complex in the nucleus [109, 117]. We detected a 100-fold suppression in the levels of transcripts expected to code for the Arp2/3 complex in infected *T. ni* hosts. Expression of a couple of transcripts coding for zipper and cytoplasmic myosin light chain proteins, also known for their roles in regulating cell shape, was reduced by over 1800-fold in the infected *T. ni* hemolymph (Supplementary Table 4). Viral infections are known to suppress host cell apoptosis as a counter defense mechanism to promote viral replication [109]. The viral gene *Ac135* is one such gene known to suppress apoptosis. *Ac135* was abundant (in the 38% highly expressed viral transcripts) in both infected hosts in our study (Supplementary Table 6). We found reciprocal coordinated suppression of several host transcripts associated with apoptosis in infected hosts. For example*, calreticulin* (*Calr*) [118]*, GDP dissociation inhibitor* (*Gdi*) [119], and *death-related protein* (*Drp*) [120] were coordinately suppressed in infected *T. ni* hemolymph (Supplementary Table 4). Notably, the characteristic host apoptosis marker genes known for their defense were absent in the transcripts identified as significantly induced in the infected hosts. Therefore, we see a bias in the host transcriptomic signals towards an overall suppression of host apoptosis as a counterdefense mechanism, favoring the budded virus propagation (Supplementary Table 4).

AcMNPV induced developmental arrest in the host is a known outcome in infected instars. In support of this expectation, we observed multiple host transcripts associated with larval developmental arrest. For example, the insect juvenile hormone synthesis genes, *adenosylhomocysteinase* and *farnesyl pyrophosphate synthase* [121], and transcripts encoding the heme peroxidase, *Cysu,* required during wing maturation [122], were co-suppressed in the infected *S. frugiperda* and *T. ni* hemolymph (Supplementary Table 4). Similarly, *Ac15*, a highly abundant viral gene in infected hosts (RPKM of 295.8 in *S. frugiperda* and 433.1 in *T. ni*, Fig. 5b and Supplementary Table 6) codes for the EGT enzyme that inactivates the insect molting hormone, ecdysone that would lead to host developmental arrest [123].

## Discussion

### Host transcriptomic signatures suggest impaired membrane integrity enabling viral proliferation

In Figure 6 we provide an overview of genes and pathways affected by host-viral interactions in the hemocoel at the systemic infection stage based on the collective deduction of our transcriptome-based analyses. Our results provide a compelling set of transcriptomic signals to support suppression of chitin-associated processes in the infected hosts, which can be linked to weakened membrane stability, as well as disrupted tracheal development during the systemic infection phase (Fig. 3, 4, and 6). Chitin-centric processes are fundamental to the transcriptional regulation that play a key role in integrating various metabolic processes operating at the cell, organ, and organism levels during pathogenesis. AcMNPV infection via occlusion derived virus is regulated by the chitin based peritrophic matrix permeability to virions in the midgut epithelium. The midgut epithelium tissue and the adjacent hemolymph in contact with the tracheal system form the focal point for systemic infections by the budded virus [5, 124, 125]. Therefore, analyzing the transcriptional profile associated with chitin in the host during host-pathogen interactions as suggested by He et al., (2020) would be an important step in studying the possibility of both using the pathogen and enhancing the virulence of the pathogen for use as a bioinsecticide.

**Figure 6:**
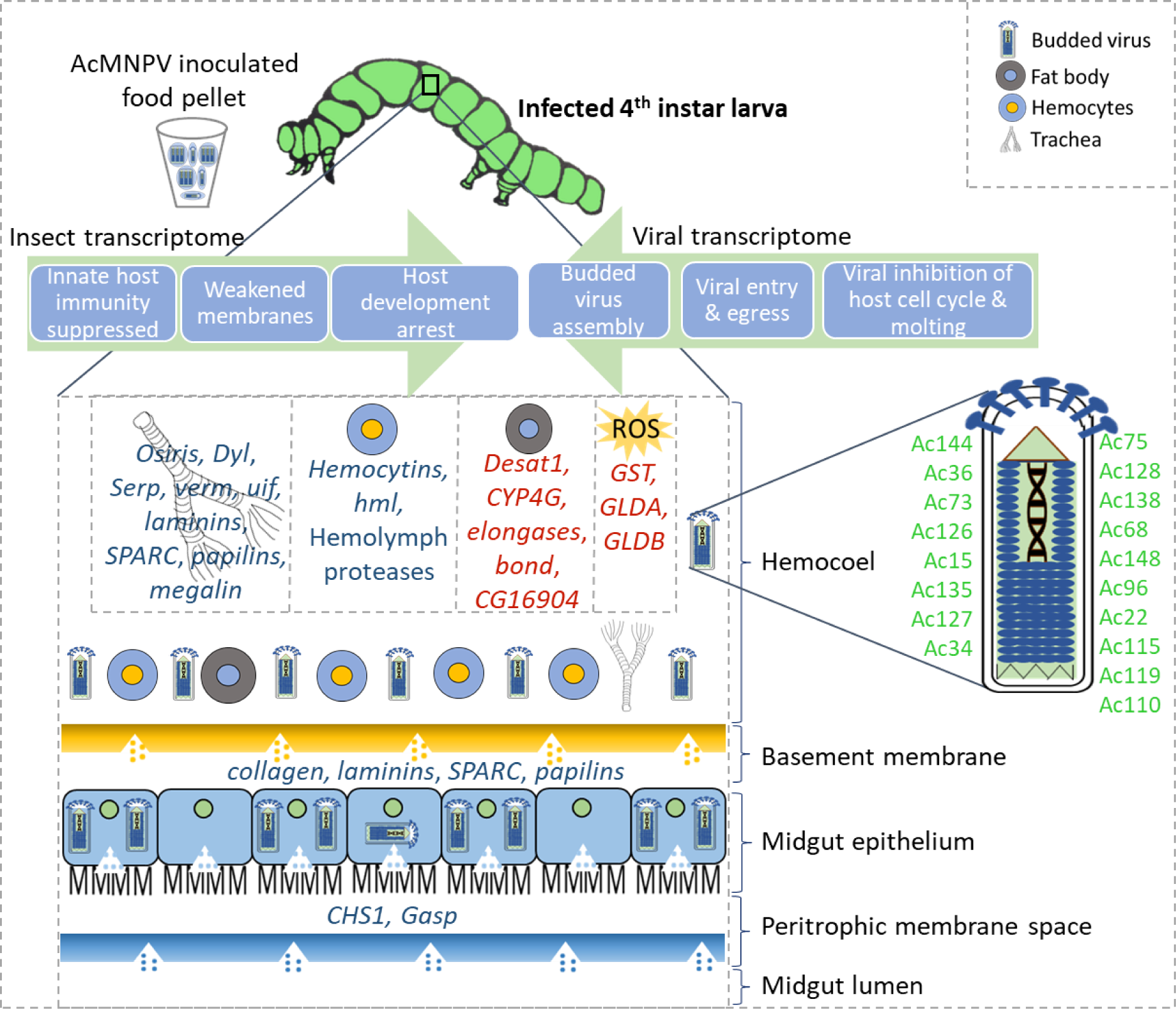
Overview of host and viral transcriptome responses in the hemocoel from a 4^th^ instar larva infected with AcMNPV at the systemic infection stage. Prominent host genes that respond to the viral infection are listed in the cells/tissues most likely to express those genes. Induced genes are in red and repressed genes are in blue. Highly abundant viral genes are given in green.

Chitinases degrade insoluble polysaccharides into soluble oligosaccharides during the molting process of insects and play indispensable roles in organ morphogenesis, cell division, and development [124, 126]. Pathogens influence host transcription of chitinases and associated proteins [127, 128] and can interfere with molting of the insect hosts [129]. However, in our current study, we did not see a significant suppression of host chitinases. Instead we found a transcript coding for a putative chitinase in *T. ni* to be significantly induced in the infected hemolymph (Fig. 4a). The AcMNPV genome also codes for a chitinase that disrupt the cuticle and peritrophic matrix of the insect host [130]. Chitinases coded by baculovirus genomes have a greater sequence similarity to bacterial chitinases involved in fungal chitin degradation and are distinct from insect chitinases both in sequence as well as localization in host tissues [107, 131]. A functional viral chitinase is critical to complete the infection cycle of the AcMNPV. In our study, the AcMNPV chitinase gene, *Ac126*, is highly expressed in both infected hosts (Fig. 5a, b, and Supplementary Fig. 6c). It is possible that the viral chitinase transcripts, together with the *cathepsin* transcripts required for liquefaction of the host, are transcribed early on during budded virus production, but are kept inactive until a later stage when occlusion bodies are produced toward the completion of the infection cycle.

The stability of basement membranes in the host is critical in mounting an innate structural barrier against the movement of the virus and containing the infection. Transcripts associated with the major glycoproteins (collagen, laminin, osteonectin, and papilin) [64–66], known to function in basement membrane stability, were all coordinately suppressed in both infected hosts in our study (Fig. 3, 4b, 6, and Supplementary Table 4). Laminin and type IV collagen are the dominant glycoproteins in the basement membrane and form a stable scaffold for other glycoproteins to create a network that provides both structural and signaling support to adjacent tissues [67–69, 132, 133]. Glycoproteins such as osteonectin bind with Ca^2+^ in the extracellular matrix and mediate cellular interactions with the matrix. These glycoproteins are required for membrane assembly, and facilitate tissue remodeling after damage to the membrane [63, 64]. They are found in fat bodies, basal lamina in the basement membrane, and in the extracellular layer secreted by epithelial cells and tracheal cells [60, 66]. Viral proteases especially target the laminins in the basal lamina of tracheal cells, making them more susceptible to virus movement, and thereby facilitate systemic infections [134]. Damage to basement membranes are unavoidable during the systemic infection of the budded virus. Therefore, the coordinated down-regulation of multiple transcripts coding for both stable and dynamic components of the basement membranes (>15% of DETs) suggests weakened barriers in the gut epithelium, hemocytes, and tracheal cells in the host. The coordinated and targeted suppression of host basement membrane proteins could be under the regulation of the viral genome in order to facilitate membrane disruption during pathogenesis in susceptible hosts.

Massive reorganization of lipid membranes is expected as the virus escapes from midgut to the hemolymph or from the hemolymph to tracheoblasts [135, 136]. A recent study by Li et al., (2018) demonstrated that fatty acid biosynthesis was induced at early disease stages and led to the reduction of virions in *S. frugiperda* Sf9 cell cultures, possibly as a host defense response. This supports the proposal that fatty acid synthesis is a key process that modulates viral infection levels in host cells [138]. In our study, we observed transcripts involved in fatty acid modifications strongly induced in both hosts in response to the AcMNPV infection (Fig. 4a and 6). While induced host transcripts were much fewer compared to the suppressed transcripts (Fig. 2a and b), it is notable that *Desat1*, stearoyl CoA desaturase, *elongases* (*bond*, *CG31523*, *CG16904*), and transcripts potentially coding for cytochrome P450 (*CYP4G1*) that collectively function in lipid biosynthesis, were among the few and most induced transcripts in the infected hosts (Fig. 4a). Taken both hosts together, lipid metabolism accounts for 50% of all induced DETs that were annotated with a known function (Supplementary Table 4).

*Desat1*, a key gene in unsaturated fatty acid biosynthesis, was also among the most induced genes in the tobacco budworm (*Heliothis virescens*) hemocytes infected with *Helicoverpa zea single nucleopolyhedrovirus* [139]. Although it is associated with starvation induced autophagy in Drosophila [140, 141], many other integral components of the autophagy pathway known to be under transcriptional regulation [142] were not noticeably impacted in our study.

Whether host lipid synthesis genes are primarily involved in disease susceptibility or resistance is not clear. Distinguishing the specific involvement of these genes is challenging partly because of inadequate functional characterizations available for many of these genes in insect hosts. For example, in line with our results, previous studies have shown that *CG16904* is induced during parasitic infections [143], but its function is unknown. Similarly, *CYP4G1*, a cytochrome P450 gene involved in cuticular lipid synthesis and highly conserved in insects, has been identified as the most highly expressed among 85 of *CYP*450 genes of Drosophila [83]. Yet, the role of CYP4G1 during viral infections has not been elucidated, despite its direct functional association with the cuticle development. It is unclear how the host defenses lead to the up-regulation of these transcripts associated with lipid synthesis specifically during viral infections concurrently to the suppression of chitin-based processes and other structural components of the basement membrane. Based on the current study from intact infected hosts and supported by previous cell culture studies, it is imperative that the specific role of lipid synthesis in the complex host-pathogen interactions during AcMNPV infection are comprehensively investigated.

### Hemocyte-mediated innate immunity is suppressed during the budded virus infection

Hemolymph is the primary target tissue we used to deduce biological processes affected by the budded virus that is known to largely invade the hemocoel. Hemocytes are known to elicit innate immune responses upon pathogen infections. During an infection, pathogens can be phagocytosed by hemocytes, agglutinated by hemolectins and other associated proteins in hemostasis, subjected to the melanization defense response triggered by hemocytes, or destroyed by oxidants or other antimicrobial compounds produced by hemocytes [144]. In our study with the AcMNPV budded virus infection progressing into a systemic infection during the 4^th^ instar larvae, we see prominent transcript signals that suggest a suppression of the hemocyte mediated immune responses rather than transcriptional induction of those primary genes involved. This inference is supported by the coordinated suppression of *hemolectin* and *hemocytin* transcripts in infected *S. frugiperda* and *T. ni* hosts together with other transcripts such as the von Willebrand clotting factor (Fig. 3, 4b, 6, and Supplementary Table 4). The larval stage specific clotting factor, *Hml* and its homolog *hemocytin* are critical genes associated with hemostasis in insects [73–75].

Serine proteases and serine protease inhibitors play vital roles in hemocyte driven phagocytosis, melanization, and antiviral immune responses in addition to their other pleotropic functions in insect development [145, 146]. The melanization reaction is tightly coupled to hemostasis reactions induced by hosts under pathogen infections as an integral part of the host immune response [70–72]. Lepidopteran hosts are known to use serine proteases produced in hemocytes to trigger melanization reactions in the hemolymph [77, 147]. Yet, the detailed functional mechanisms of specific serine proteases in mounting defense responses against baculoviruses are poorly understood. In our study, a number of serine proteases and serine protease inhibitors were co-suppressed in both infected hosts (Fig. 4b and Supplementary Table 4), implying a defense response compromised by the virus in infected hosts. It should be noted that both *S. frugiperda* and *T. ni* are known to be highly permissive hosts to AcMNPV infections [148, 149].

The lepidopteran innate immunity elicited by hemocyte aggregation and hemolymph melanization against bacterial pathogens is well established [70–72]. However, their role is largely unexplored under baculovirus infections, partly because, most studies have used cell lines and not intact tissues. Based on our results, the presence of the budded virus appears to strongly suppress the host immune responses initiated via hemocytes.

### During disease progression, cellular energy usage is altered with substantial consequences in redox homeostasis, primary metabolism, and development of the entire organism

Synergistic to maintaining membrane integrity via coordination of chitin and lipid metabolism, host cell survival depends on being able to maintain energy metabolism and redox homeostasis to minimize oxidative stress during infection and prevent further damage to membranes and DNA [150]. The complex regulation of energy metabolism is tightly coupled to the cellular redox state and plays a central role in viral infections. Therefore, a failure to maintain homeostasis of these critical pathways suggests early signs of systemic progression of infection.

We found multiple transcripts potentially coding for integral enzymes in primary energy metabolism and redox homeostasis suppressed in both host species by AcMNPV infection (Supplementary Table 4). Lavington et al., (2014) have demostrated that a handful of enzymes in the central energy metabolism can shift the flux balance and energy homeostasis. These are often found to be regulated at the transcriptional levels. Due to their high connectivity to many primary metabolic pathways, the transcripts of these flux-controlling enzymes can be used to sense the energy state of the cell. One such key enzyme in maintaining the redox pools and energy balance is the malic enzyme (coded by *Men* and *Men-b* genes) that catalyzes malate to pyruvate while reducing NADP to NADPH [151, 152]. It has been estimated that 30% of the total cytosolic NADPH is produced by *Men* in Drosophila [153, 154] and it is a critical enzyme in coupling energy metabolism to ROS levels under oxidative stress. Transcripts coding for the malic enzyme were suppressed with several other glycolytic transcripts in *T. ni*, suggesting a transcriptional signal of altered redox balance in this host. Redox imbalances can cause severe oxidative stress leading to cell fatality. Specially for viral pathogens, host defense responses primarily include oxidative stress mitigation and ROS scavenging [155, 156]. Notably, three out of the four significantly induced transcripts in *S. frugiperda* (Fig. 4a, 6, and Supplementary Table 4) are transcripts that code for enzymes that are induced as a defense response to minimize oxidative stress [85–87]. To further connect such components into biological pathways and identity specific molecular targets during baculovirus infections, a critical mass of genetic studies needs to accumulate on specific gene functions and transcriptomic responses in multiple lepidopterans.

The overall transcriptomic profiles in both infected hosts also suggest a compromised or reduced allocation of energy into other critical larval development processes. We identified a number of transcripts critical for the development of wings, muscles, renal functions, and neurons in both infected hosts significantly suppressed (Supplementary Table 4). Concurrently, we see a substantial fraction of ribosomal protein transcripts down-regulated in infected *T. ni* implying altered rates for protein translation and overall metabolism (Supplementary Table 4). This reduction is also observed in *S. frugiperda* but to a lesser magnitude. A number of ribosomal proteins were down-regulated in *S. frugiperda* in response to the AcMNPV infection at a significant level (q-value ≥0.95), but the fold change was marginal (less than 4-fold) (Supplementary Table 4). Taken together, these results suggest that critical cellular and metabolic processes seem to have been significantly affected, even if only 1% of the transcriptome in the infected hosts showed significant reduction in response to the AcMNPV infection. The impaired cellular and metabolic processes consequently may have affected insect development as suggested by the suppression of several transcripts associated with the insect juvenile hormone synthesis and molting hormone regulation. In summary, AcMNPV infection affects multiple processes from cellular to whole organism level.

### Signaling processes associated with AcMNPV infection

The infected hemocoel of both hosts is expected to carry disease signaling as a systemic signal to activate immune responses as well as signaling through pheromonal pathways. Pheromonal signaling in insects is widely studied as a form of chemical signaling that can lead to aggregation of individuals specially during reproduction [157]. Several studies have discovered long-chain fatty acids that attract other larvae using novel pheromonal signaling pathways as a mechanism during immature larval stages to aggregate individuals [158–162]. While pheromonal signaling involves complex genetic and metabolic networks, Desaturase1 (Desat1) is a key enzyme that is associated with pleiotropic effects on both pheromone production and perception [163–165]. Similarly, the lipid elongase gene, *bond* is also required for pheromonal signaling and known for its role in conspecific signaling [68]. *Desat1* and *bond* are co-induced in infected hosts (Fig. 4a and 6), while, fatty acid biosynthesis and pheromone metabolism were among the enriched functions in response to the AcMNPV infection in our study (Fig. 6 and Supplementary Fig. 4). The underlying genetic mechanisms of how pheromonal signaling pathways may have been exapted into a disease signaling pathway is unknown, but previous studies have confirmed the induction of these pathways in insects during viral infections [162, 166, 167]. The induction of a pheromonal pathway leading to conspecific aggregation during baculovirus infections could facilitate disease progression between individuals as non-infected larvae in close proximity to larvae that are undergoing liquefaction have a high risk in getting infected in the next disease cycle. Therefore, a pleiotropic gene such as *Desat1* is a likely candidate to be co-opted for behavioral traits evolved under an arms race between baculoviruses and their lepidopteran hosts.

Alternatively, lipid synthesis genes could play a role in disease signaling systemically within the infected larvae by triggering ROS signaling [141, 168, 169]. The co-induction we observed for *GST* and other oxidative stress indicators (Fig. 4a and 6) in *S. frugiperda* may further support this idea of the involvement of ROS pathways in disease signaling.

### The role of AcMNPV genes found in the host hemolymph

The AcMNPV protein-coding genes regulate host cellular and physiological processes as well as the production of the two distinct types of enveloped virions: the occlusion-derived virions and the budded virions [15]. Our viral transcript quantification suggests that the budded virions are more abundant than occlusion-derived virions in the infected hemolymph samples (Fig. 5a and Supplementary Fig. 6a and b), an observation also supported by previous studies [5]. The occlusion-derived virion is primarily involved in the individual-to-individual transmission, while the budded virion is used for cell-to-cell transmission within an individual.

The transcriptomic signature of the eukaryotic host genome is overrepresented compared to the viral genome expressed in our RNA samples that capture the host-parasite interactions. Yet, the specific quantification of transcripts made feasible with RNA-seq data allows the detection of clear biological signals from the viral parasite in host tissue. The infected 4^th^ instar individuals of *T. ni* had a higher proportion of viral transcripts per million reads sequenced as well as a lower viral dose needed to achieve LD95 compared to *S. fruigiperda* (Fig. 1, 5a, and Supplementary Table 2b).

### Viral entry to cells, assembly, and egress

The two baculovirus virion types have both distinct and shared nucleocapsid and envelope proteins that serve as structural components, and perform roles in entry and exit from cells [5]. The budded virions following their initial budding from the midgut epithelial cells get circulated in the hemolymph where they can bind and enter most cell types in contact with the hemolymph [5, 149]. Many of the essential genes that function in the egress pathways also tend to have functions in forming the nucleocapsid or the envelope. For example, *Ac75* is a core gene that is required for exiting the nucleus in the egress pathway used by the budded virus, and it is involved in the formation of intranuclear microvesicles as well as envelope and nucleocapsids of the occlusion-derived virus [170, 171]. It is found to be the second most highly expressed viral gene in both hosts in our study (Fig. 5b and Supplementary Table 6).

The entry of occlusion-derived virus into the midgut epithelial cells primarily depends on a protein complex formed of nine core PIF proteins that are integral to the occlusion derived envelope [5]. It is interesting to note that eight of these nine viral transcripts, *PIF-0/Ac138, PIF-1/Ac119, PIF-2/Ac22, PIF-3/Ac115, PIF-4/Ac96, PIF-5/Ac148, PIF-6/Ac68, PIF-7/Ac110* were not only detected in the infected hemolymph samples in our study, but also found at a very high expression level ranging from 101 to 728 RPKM in infected samples (Fig. 5a and b). It is unclear why we observed such a striking signal for *PIFs* in the hemolymph that could be associated with the occlusion derived virus. Those samples with any extraneous tissue such as midgut residue were not used for further processing to avoid contamination of our hemolymph samples used for RNA extraction. It is possible that these PIF genes are transcribed but not translated until much later or PIF proteins may have yet-to-be discovered roles in the budded virus stage.

The cell recognition and entry of budded virus into the host cells is primarily controlled by a single glycoprotein, GP64 coded by the core gene, *Ac128* [5]. This is also one of the most abundant envelope proteins in the budded virus that functions in binding with the host plasma membrane [172–174]. As expected, we detected very high transcript abundance for *Ac128* in both infected hosts (Fig. 5b, RPKM of 461 in *S. frugiperda* and 937 in *T. ni*). Given that *Ac128* is easily detectable in both infected hosts; its essential role in viral entry into cells that can initiate systemic and irreversible infections leading to the death of the host; and its high sequence conservation [175, 176], make it an attractive candidate gene in the search for molecular targets best suited to create host specific biopesticide developemnt.

### Viral genes that influence host cell cycle and molting

The host cell cycle regulation affected by the viral genes is among the most invariable processes expected during host-viral interactions. AcMNPV is known to cause cell cycle arrest in their lepidopteran hosts [177]. Therefore, in our study it was not surprising to detect a major cell cycle inhibitor, Ac144/odv-ec27 coding for a cyclin as the highest expressed virus gene in both host species (Fig. 5a) [19, 104]. A large number of cyclins serve as key checkpoint regulators in the complex gene regulatory network of the eukaryotic cell cycle [178]. Therefore, future studies investigating specific gene-to-gene targets of host cyclins and their viral cyclin inhibitors could identify viral strains targeting a specific host or even a specific developmental stage of the host to facilitate safer biocontrol using baculoviruses.

Baculoviruses arrest the molting of infected lepidopteran larvae [15, 91]. This process is primarily governed by the ecdysteroid UDP-glucosyltransferase (EGT), a viral enzyme that inactivates the insect molting hormone, ecdysone [123]. *Ac15* in the AcMNPV genome codes for EGT. In our study, *Ac15* is among the most highly expressed viral transcripts found in the infected hemolymph of *S. frugiperda* and *T. ni* (Fig. 5b, 6, and Supplementary Table 6). Previous studies also have reported higher EGT activities in the hemolymph compared to other tissue [179]. There is great interest in the viral induced behavioral effects of lepidopteran hosts since Hoover et al., demonstrated the role of EGT on the climbing behavior of gypsy moth larvae [180]. Subsequent studies have confirmed the role of EGT in influencing behavioral traits of other hosts including *T. ni* and *Spodoptera exigua* [181, 182]. However, much of the genetic basis is unknown for these behavioral traits and at least in *Spodoptera* hosts, EGT alone is reported to be insufficient to elicit behavioral traits [181].

### Host transcriptional responses to the budded virus during the systemic infection stage differs from the midgut responses to the occlusion-derived virus during the primary infection stage

Shrestha et al., (2019) described the host transcriptomic landscape of the midgut during the primary infection phase of AcMNPV in *T. ni* 5^th^ instar larvae primarily caused by the occlusion-derived virus. The current study that focusses on the systemic infection stage predominantly caused by the budded virus in 4^th^ instar larvae of two lepidopteran hosts including *T. ni* depicts a very different host transcriptomic landscape. The most consistently up-regulated transcripts (at least 16 fold) observed in the midgut in the study by Shrestha et al., (2019) included, *REPAT* (*REsponse to PAThogens*), *Atlastin* (involved in ER and vesicle trafficking), cyclic GMP-AMP synthase (*cGAS*) genes that can bind to cytosolic viral DNA*, 3 ubiquitin ligase SIAH, a zinc finger CCHC,* a *peroxidase,* and a *chymotrypsin-like serine protease.* None of these transcripts were found to be significantly expressed in response to the infection in the infected hemolymph in our study. An earlier study had shown increased *REPAT* in the midgut of baculovirus infections of *Spodoptera exigua* larvae [183] similar to the observations made by Shrestha et al., (2019) for *T. ni.* These previous observations and the absence of significant changes to these transcripts in the hemolymph during systemic infections imply that these host transcripts may be specific to the infection phase or tissue. There is a clear transcriptomic signal given by multiple key apoptosis-related genes induced in the infected midgut of *T. ni* as reported by Shrestha et al., (2019). While we do not observe the induction of the same transcripts in our study, several other apoptosis-related genes were suppressed in the infected hemolymph during systemic infections (Fig. 6).

For certain time points post-infection in the midgut, Shrestha et al., (2019) reported up-regulation for several cuticle-related transcripts. The transcriptomic signal associated with cuticle-proteins are likely stemming from tracheaoblasts in the hemolymph in our study contrasting to the transcripts reported by Shrestha et al., (2019) likely coding for cuticle-proteins affected in the peritrophic matrix lining the midgut during the occlusion-derived virus propagation. The invasion of the budded virus into the tracheal epidermis is essential to the progression of the systemic infection as the host cannot shed these cells unlike the gut epithelium infected by the occlusion-derived virus that can be shed as seen in semi permissive hosts [149]. This may explain why we observe exceedingly more transcripts potentially coding for cuticle, chitin, and associated membrane processes clearly suppressed as a result of successful disease progression than in infected midgut cells reported by Shrestha et al., (2019).

The most notable consistently down-regulated genes (by at least 16-fold), during the occlusion-derived virus invasion of the midgut, mainly included orthologs of *flippase,* and genes coding for a number of Cytochrome P450 enzymes, serine proteases, calcium binding protein P, and dehydroecdysone 3 alpha-reductase as noted by Shrestha et al., (2019). None of these were significantly changed during the budded virus infection in either host in the current study. While there were hardly any direct overlap of down-regulated transcripts between the primary-midgut infection and the secondary-systemic infection, we see melanization as a suppressed pathway in both studies. Shrestha et al., (2019) described the down-regulation of *serine proteases* involved in the melanazation cascade similar to our observation for the suppression of multiple serine proteases thought to be involved in melanization and other defense responses (Fig. 5b). Contrasting to the overall observations made by Shrestha et al., (2019) during the midgut infection, the hemolymph of both hosts in our study during systemic infection appear to clearly induce transcripts associated with oxidative stress while suppressing those related to hemostatis, chitin metabolism, and tracheal development.

### Key host genes affected by the AcPNMV infection are targets of commercially available pesticides used against lepidopteran pests

The baculovirus genes directly regulate primary metabolic pathways of the host during viral replication that overwhelms the energy balance of host cells, eventually leading to cell death. The commonly targeted host genes by the viral pathogen include *CHS1*, and transcripts associated with actin driven cellular functions as well as genes involved in insect hormone regulation. It is interesting to note that many of the chemical insecticides also use the same genes as primary targets to control lepidopteran pests. However, unlike chemical insecticides, baculoviruses continue to spread in the field post-host liquification.

Many insecticides developed against insect pests target chitin biosynthesis as a more specific and safer alternative to generic insecticides such as pyrethroids and organophosphates. These chitin synthesis inhibitors largely include the benzoylphenylurea (BPU) group of insecticides, oxazolines, tetrazines, thiadiazines, thiazolidines, and triazines [184, 185]. All chitin biosynthesis inhibitors act on chitin synthesis at various stages of the complex biochemical pathways leading to the interruption of chitin production and cuticle development. The BPUs are shown to target *CHS1* to inhibit chitin metabolism early in the biosynthesis pathway [186]. Notably, *CHS1/kkv* is the main chitin synthase required for epicuticular stability, intact procuticle, maintenance of epidermal morphology, and sclerotization and pigmentation of the cuticle [187]. A number of genes associated with chitin synthesis and cuticle modifications (discussed earlier) are among the most highly suppressed transcript cluster in both hosts during the systemic infection.

Pyridalyl is a commonly used potent insecticide against lepidopteran pests [186]. It has been used to control fall armyworm outbreaks in South Africa [188, 189]. The molecular mechanism of Pyridalyl generates excessive amounts of ROS that eventually leads to severe oxidative stress and cell death in lepidopterans [190]. Among the handful of strongly induced genes during the systemic infections of the budded virus, *GST* and other genes associated with oxidative stress are notable (Fig. 4a and Fig. 6). Further induction of these oxidative stress pathways disproportionately divert energy to oxidative stress responses that could expedite cell death and, in turn, host death. The current observation made in our study further supports the insecticidal potential of AcMNPV strains selected to induce host oxidative stress responses similar to what observed with the Pyridalyl activity.

Double stranded RNAs (dsRNAs) that mimic insect transcripts have emerged as a powerful tool for targeted pest control. For example, dsRNAs of actin transcripts used as foliar sprays have shown to be a promising insecticide for Colorado potato beetles that damage multiple Solanaceae crops [191]. In line with our findings made in the current study, baculoviruses are known to target actin-mediated cellular processes. Actin is present in all cells and customization to target-specific lepidopteran actins or a regulatory gene of actin-mediated processes is equally achievable with baculoviruses. Further, baculoviruses are more effective as delivery agents in controlling host genes than the passive delivery methods available for dsRNA-based insecticides [192]. The use of recombinant baculovirus strains to control pests has been proposed for over decades and has recently gained more attention as sustainable biopesticides [193–196]. Transcript level inhibition of the juvenile hormone biosynthesis or alterations to its regulation is a common target attempted in recombinant baculoviruses developed as potential biopesticides [197, 198]. Host genes associated with juvenile hormone regulation were noticeable among suppressed transcripts specially in the infected *T. ni* even when wild type AcMNPV strains were used [199] similar to observations made in our current study.

## Conclusions

We identified extensive overlap between biological processes that were represented by differently expressed genes in the two hosts in response to the virus as well as convergence of functional clusters of genes expressed in the virus in response to the two hosts. The overall host transcriptomic signals suggested chitin-associated processes and basement membrane integrity were compromised together with hemocyte-initiated immune responses in both infected hosts. Oxidative stress indicators, moderately induced by the viral infection, may play a role in systemic disease signaling with the induction of selected classes of fatty acids (Fig. 6). The entire core viral genome was expressed during the systemic infection phase in both hosts, with a bias towards processes associated with budded virus production and transport. The host-virus interactions deduced from co-expressed host and viral transcripts indicate an overall transcriptomic landscape overwhelmed by viral counter defenses that facilitate disease progression. The specific transcripts and the convergent biological processes, highlighted in our study as highly affected during infections, identify key genes and pathways as potential molecular targets in designing recombinant AcMNPV strains as molecular tools in sustainable pest management.

## Methods

### Insect and virus source material

Given the natural progression of an epizootic in the field and the need to collect a considerable amount of hemolymph for the transcriptome analysis, we used 4^th^ larvae in the experiments outlined below. *S. frugiperda* and *T. ni* were obtained as eggs from Benzon Research Inc. (Carlisle, PA, USA). Once the eggs hatched, we reared them in individual one-ounce cups on artificial diet (Southland Products Inc., Lake Village, AR, USA) at 28.9 °C and a 16 hour-light and 8 hour-dark cycle until they reached the 4^th^ instar. Wild-type AcMNPV strain E2, which was used in this study, was field collected. To amplify the virus for the experiment, the virus was passed through *Chrysodeixis includens*, the soybean looper.

### Determination of LD95 of AcMNPV for *S. frugiperda* and *T. ni*

We used a standard dose-response protocol and Bayesian analysis to quantify the lethal dose at which 95% of the larvae would be expected to succumb to viral infection or the LD95. For the experiment, thirty recently molted 4^th^ larvae, which were starved for 24 hours, were fed a known amount of virus on a small diet cube. The virus was suspended in a 3 μl droplet of deionized water. One set of larvae was used as a control and consumed a diet cube that had been only inoculated with 3 μls of deionized water. None of the controls became infected. Only larvae that consumed the entire diet cube were used in the experiment to ensure that the larvae received a full dose of the virus. Viral doses varied depending upon the species (Fig. 1). After consuming the diet cube, larvae were placed on one-ounce cups and reared until pupation or death. Death resulting from AcMNPV infection was confirmed either by host liquefaction in the diet cup or by examining hemolymph under a light microscope [7]. For *T. ni*, the experiment was conducted twice, since the first set of experiments used doses that were too high resulting in almost 100% mortality and, thus, making it difficult to estimate the LD95. The second set of experiments used much lower doses. We combined the data from the two experiments for the *T. ni* dose-response analysis.

To analyze the data, we used a Bayesian framework with vague priors to fit a logistic regression model [200] for each species. The associated slope and intercept of the fitted model was used to calculate the LD95. All analyses were conducted in R (R core Team, 2018) using the JAGS [202] and the R2JAGS packages [203]. For each analysis, three chains were run from different starting points. The first 10,000 draws from the Bayesian Markov chain Monte Carlo (MCMC) were removed to account for transient dynamics at the start of the chain. The remaining 90,000 draws were retained to estimate the parameters of the logistic regression. All non-discarded draws were retained to ensure precise parameter estimates [204]. After a visual inspection of the chains for convergence, multiple tests were used to ensure that the chains had converged including the Gelman-Rubin and the Hiedelberg-Welch tests [205]. The chains for each analysis were combined to form a posterior distribution. Additionally, we conducted a posterior predictive check to test whether the predicted model fit the data collected [206]. As part of the posterior predictive check, Bayesian p-values were calculated. Values near 0.50 indicate that the model fits the data reasonably well [207]. The Bayesian logistic regression for both species passed each of the individual tests outlined above.

### Insect treatment with AcMNPV virus

Using the LD95 calculated from the dose-response experiments, 4^th^ instar larvae from both species were fed the appropriated dose of virus (*S. frugiperda*, 10^4.5^ OBs; *T. ni*, 10^3^ OBs) on a diet cube using the same method as the dose-response experiment. Control larvae were fed a diet cube inoculated with deionized water. After 30 hours, 30 individuals per sample were used to extract the hemolymph.

### Extraction of hemolymph total RNA and preparation of RNA-seq libraries

Prior to hemolymph extraction, each individual was chilled to ease the extraction process. A pre-chilled microcentrifuge tube was filled with a 25 μl solution containing 10 units of RNAseOut in a 0.1 % PTU dissolved in a PBS solution. The rear proleg of the 4^th^ instar larva was then cut with micro scissors. We collected hemolymph from the wound and pipetted the hemolymph into a pre-chilled Eppendorf tube. The solution was then vortexed, immediately placed in a dewar filled with liquid nitrogen, and stored at −80 °C until needed.

Total RNA was isolated from hemolymph samples using RNeasy Mini Kit (Qiagen, Hilden, Germany). On-column DNase digestion was carried out with the RNase-free DNase Kit (Qiagen), followed by a further purification step using RNeasy Mini Spin Columns (Qiagen). The quantity, quality, and integrity of the total RNA was sequentially assessed using the A260/A280 values reported with a Nanodrop spectrophotometer (Thermo Scientific, Wilmington, DE), agarose gel electrophoresis, and a BioAnalyzer (Agilent Technologies, Inc.).

RNA-seq library preparation and sequencing were done at the University of Illinois at Urbana-Champaign Roy J. Carver Biotechnology Center. Ribosomal RNA (rRNA) depletion was performed on the RNA samples using the RiboZero kit (Illumina, San Diego, CA) following the manufacturer’s instructions. Capturing polyA-enriched RNA from total RNA is a more customary approach for eukaryotic RNA-seq experiments. However, we decided to use rRNA depleted samples because we planned to identify both insect and viral transcripts which may not always contain 3′polyA sequences. The rRNA-depleted samples were used for TruSeq Stranded RNA Sample Prep kit to produce 5′ to 3′ strand-specific cDNA libraries (Illumina). A TruSeq SBS sequencing kit version 3 (Illumina) was used following the manufacturer’s instructions to generate the sequencing libraries. All libraries were pooled, barcoded, and multiplexed on two lanes of an Illumina HiSeq2000 platform to run for 101 cycles. Randomly selected reads of 100 nucleotide lengths from each library were processed and demultiplexed with Casava 1.8.2 that generated over 370 million reads with quality scores over 30.

### Sequencing, assembly, and annotation of the reference transcriptome

To allow accurate identification of host transcripts from two species, we needed to create two reference transcriptomes for the hemolymph of 4^th^ instar caterpillars. RNA-seq reads were processed to generate a reference transcriptome assembly and annotation following a custom pipeline published previously [208]. Briefly, raw Illumina reads were subjected to quality checks using FastQC and *de novo* assembled using Trinity v2.2.0 [209] using default parameters. Contigs with low read support, contaminants, and artifacts were removed as described in Oh et al., 2015. We further clustered contigs showing >95% sequence identity over >70% of total contig length of the shorter contig in each pairwise alignment, using CD-HIT-EST v4.6 [211] to minimize redundancy. For each cluster, the transcript with the longest open reading frame (ORF), predicted by Transdecoder v2.0.1 (https://transdecoder.github.io/), was selected as a representative transcript model in the final protein-coding reference transcriptome. The completeness of each reference transcriptome assembly was evaluated using Benchmarking Universal Single-Copy Orthologs (BUSCO) database v2.1 [212] with the metazoan dataset (metazoa_odb9) and default settings. A series of sequential BLAST searches found the best possible annotation for both coding and non-coding transcript sequences, using the NCBI Drosophila mRNA database, NCBI-insects-reference RNA (refseq_rna), and NCBI-non redundant (nr) databases for all eukaryotic proteins and RNA, with a maximum e-value cutoff of 10^−5^.

An ideal transcriptome is expected to consist of all expressed genes in a given condition. This would include both coding and non-coding transcripts. However, the non-coding transcript pool is highly incomplete even for the premier model species. Therefore, it would be impractical to assign reasonable functional annotations for contigs that may represent true non-coding transcripts in our study. Additionally, without any canonical structural features to use in assessing the completeness of non-coding transcripts, those transcripts could also contain a highly fragmented fraction of the assembly. Therefore, we divided our assembled transcriptome into coding and non-coding reference transcriptomes and only used the protein-coding transcriptome for our current analyses. Despite the lack of resources to fully annotate putative non-coding transcripts, this pool of non-coding transcripts likely represents a genetic component that has potential to be useful as a collective resource from diverse species as more high throughput data driven projects are conducted. Therefore, we include Table 1 and Supplementary Fig. 1, where we report a total of 101,169 and 147,772 processed non-coding transcripts, with a mean length of 495 and 549nt, for *S. frugiperda* and *T. ni*, respectively, as an additional molecular resource included in our data deposit to NCBI BioProject PRJNA664633.

The protein-coding reference transcriptome was used for the downstream RNA-seq analysis. Each sequence used as a proxy to represent gene/transcript models in our study when assessing biological processes will be designated by its gene name, followed by the shortened form of the gene name (if available), the sequence ID given by our annotation process, and the FlyBase or NCBI accession number used for its annotation in parenthesis, as in the example, *Chitinase6* (*Cht6*, TR50740|c0_g1_i1/ FBgn0263132).

### RNA-seq analysis

The goal of our experiment was to search for shared disease responses inferred from the two host species affected by AcMNPV infection using three sets of biologically independent RNA-seq datasets. Two datasets were from *S. frugiperda* and one set was from *T. ni.* The RNA-seq reads were aligned to the relevant reference transcriptome using bowtie [213] with a seed alignment length per read set to 50nt. Reads unambiguously mapped to each gene model were counted using a custom python script to generate read-count values as a proxy for gene expression. We used NOISeq [214] with a q-value cutoff of ≥0.95 to identify transcripts differently expressed between control and AcMNPV-infected samples in both insect species. Gene ontology (GO) terms enriched among differently expressed transcripts (DETs) were detected using BiNGO at FDR adjusted p-value ≤ 0.05 [215]. We used the entire reference protein-coding transcriptomes as custom backgrounds to test for functionally enriched clusters when inferring the shared biological processes identified from each host species. GO annotation of reference protein-coding transcriptomes for the two insect species was based on sequence similarity compared to *Drosophila melanogaster* gene models that have assigned GO terms. We used GOMCL [216] to identify the non-redundant functional clusters from the primary set of enriched functions generated using BINGO [215] for each species.

To assess the transcripts originating from the viral genome, particularly in the infected samples, RNA-seq reads were mapped to the AcMNPV reference genome [16] using bowtie [213]. The read counts mapped to the viral genome were normalized by converting to RPKM values (Reads Per Kilobase Million) for each viral gene expressed in the insect transcriptomes. Total read counts were calculated by adding the reads mapped to the viral genome and insect gene models for control and AcMNPV infected samples as used in a previous study [217].

**List of abbreviations**
AcMNPV: *Autographa californica* Multiple Nucleopolyhedrovirus
BUSCO: benchmarking universal single-copy orthologs
DETs: differently expressed transcripts
DNA: deoxyribonucleic acid
FDR: false discovery rate
GO: gene ontology
ORF: open reading frame
RNA-seq: RNA sequencing
ROS: reactive oxygen species
RPKM: read per kilobase of transcript per million mapped reads
TCA cycle: tricarboxylic acid cycle

## Declarations

### Ethics approval and consent to participate

Not Applicable

### Consent for publication

Not Applicable.

### Availability of data and materials

The RNA-seq data sets from this article have been deposited to the NCBI Sequence Read Archive (SRA) Database, under the accession number for NCBI BioProject ID: PRJNA664633.

### Authors’ contributions

MD and BE designed the experiment; BE conducted the dose response test of the host species, raised larvae in control and infected groups, and extracted hemolymph samples; SC extracted RNA and optimized methods to obtain high quality RNA to be used for RNA-seq libraries; PP, DHO, and MD developed the bioinformatic analyses; PP conducted the bioinformatics analysis; PP and MD wrote the manuscript with input from all authors.

## Acknowledgements

The authors thank the High Performance Computing at Louisiana State University (HPC@LSU) for providing computational resource for data analysis, Louisiana State University Library services for aiding with retrieving articles via interlibrary loans, Forrest Dillemuth for rearing larvae, and Chathura Wijesinghege and Prava Adhikari for their art work in the illustration.

## Authors’ information

All authors: Department of Biological Sciences, Louisiana State University, Baton Rouge, LA 70803, USA.

---

**Supplementary Figure 1.**
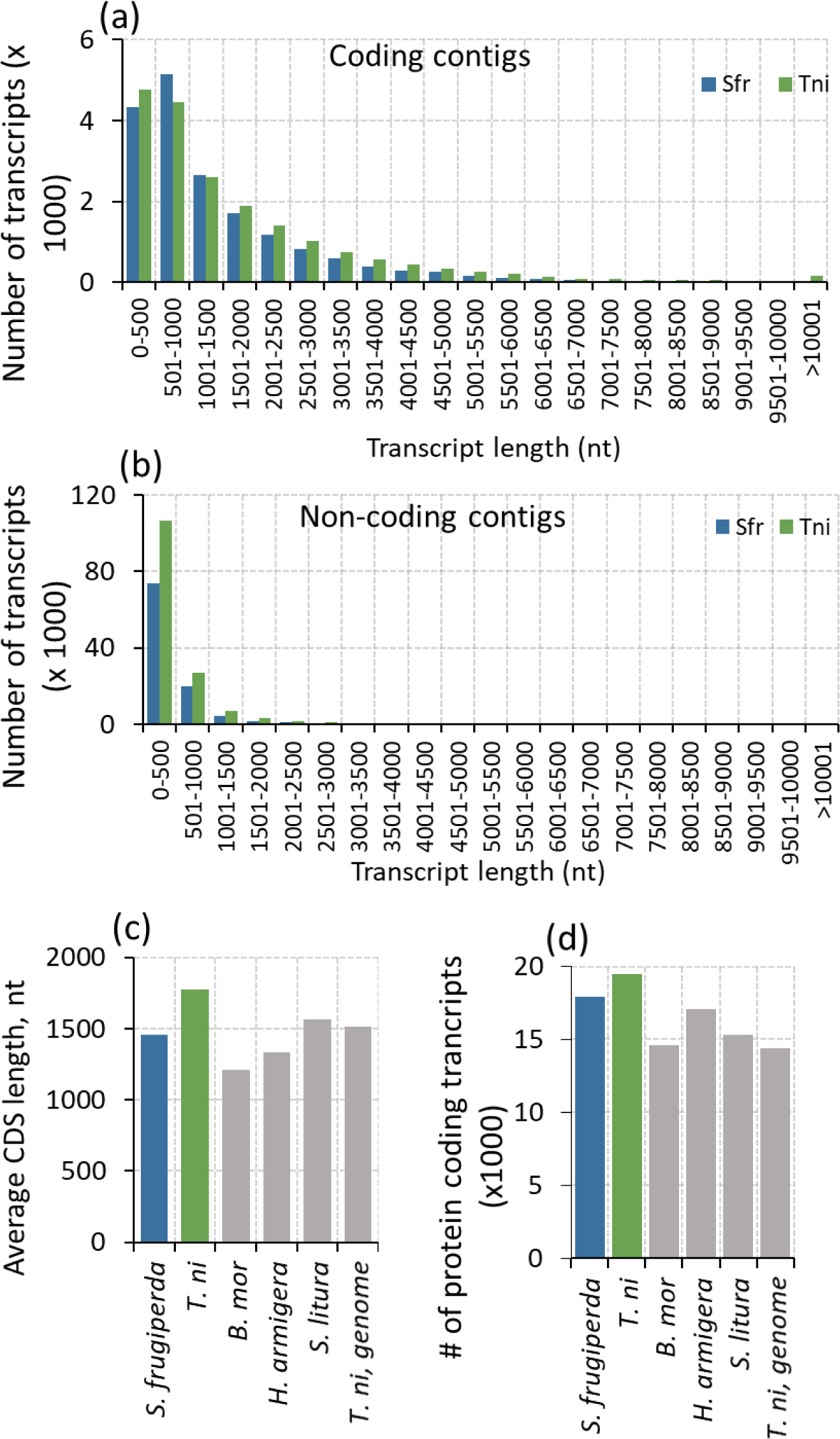
Assembled contig length frequency distribution for *S. frugiperda* and *T.ni* reference transcriptomes. [a] Coding contigs and [b] non-coding contigs. Comparison of average CDS length [c] and number of protein coding transcripts [d] between *S. frugiperda* and *T. ni* compared to *B. mori*, *H. armigera, S. litura*, and the *T. ni* genome.

**Supplementary Figure 2.**
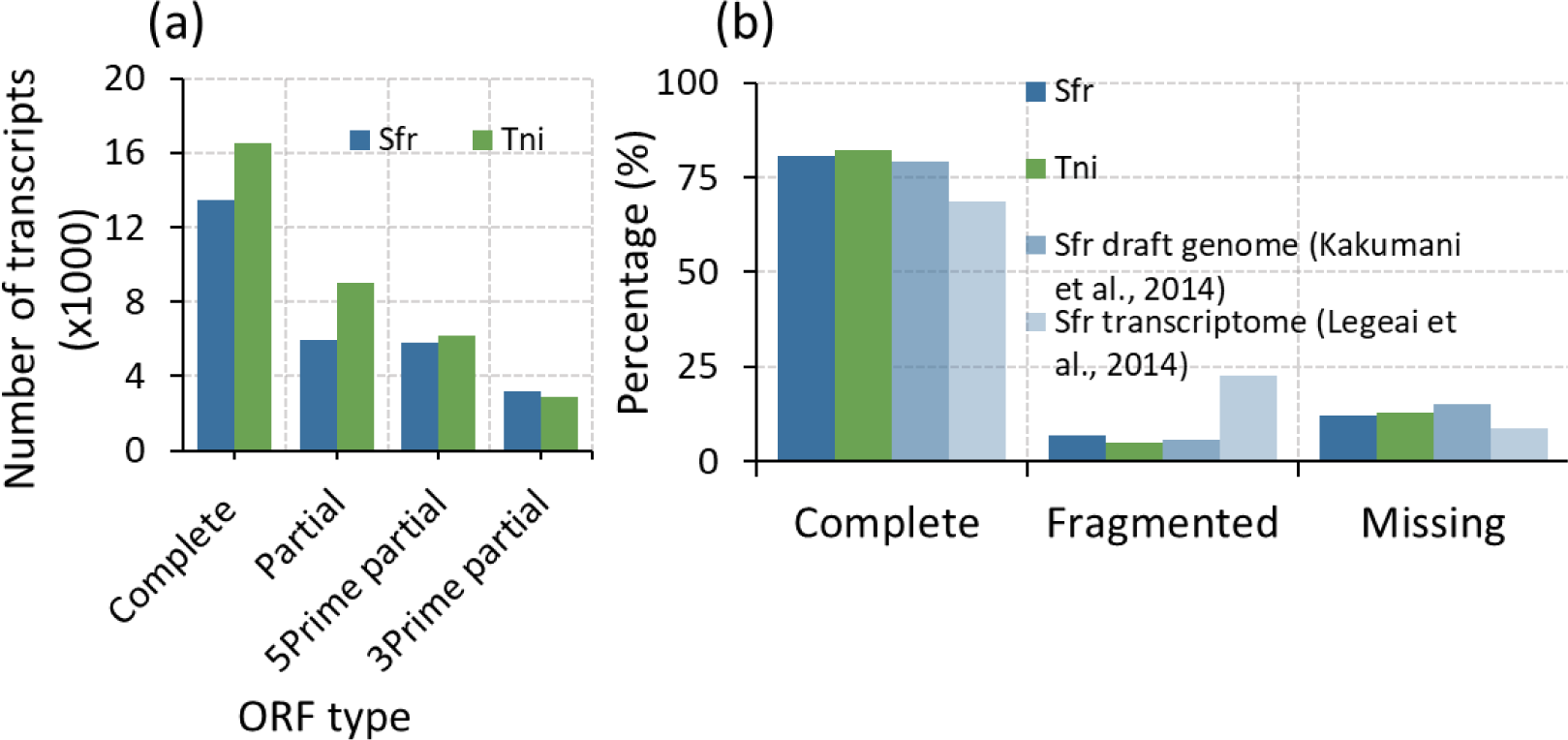
Quality assessments of *S. frugiperda* and *T. ni* reference transcriptomes. [a] proportions of different ORF types and [b] assembly completeness of the references created in this study compared to the previously published *S. frugiperda* draft genome and transcriptomes (Kakumani et al., 2014 and Legeai et al., 2014) assessed using BUSCO.

**Supplementary Figure 3.**
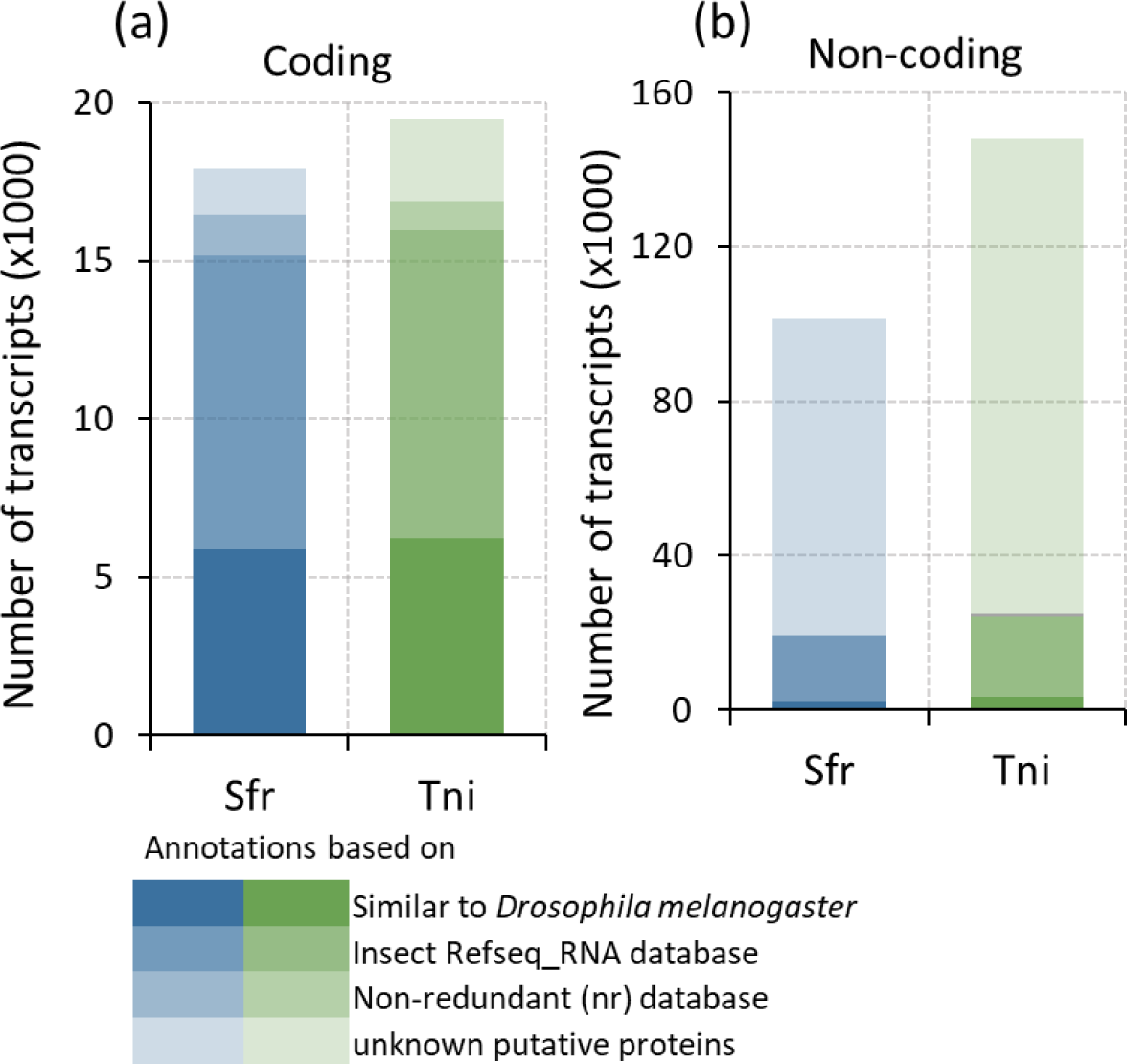
Annotation summary of the *S. frugiperda* and *T. ni* transcriptome assembly for coding transcripts. Functional annotation of reference transcriptome was performed using sequential BLAST with an e-value cutoff 10^−5^ searched within the drosophila mRNA database, insect reference RNA (refseq_rna) database, and non-redundant (nr) database. The annotations of reference transcriptome for both species are provided in the Supplementary Table 3.

**Supplementary Figure 4.**
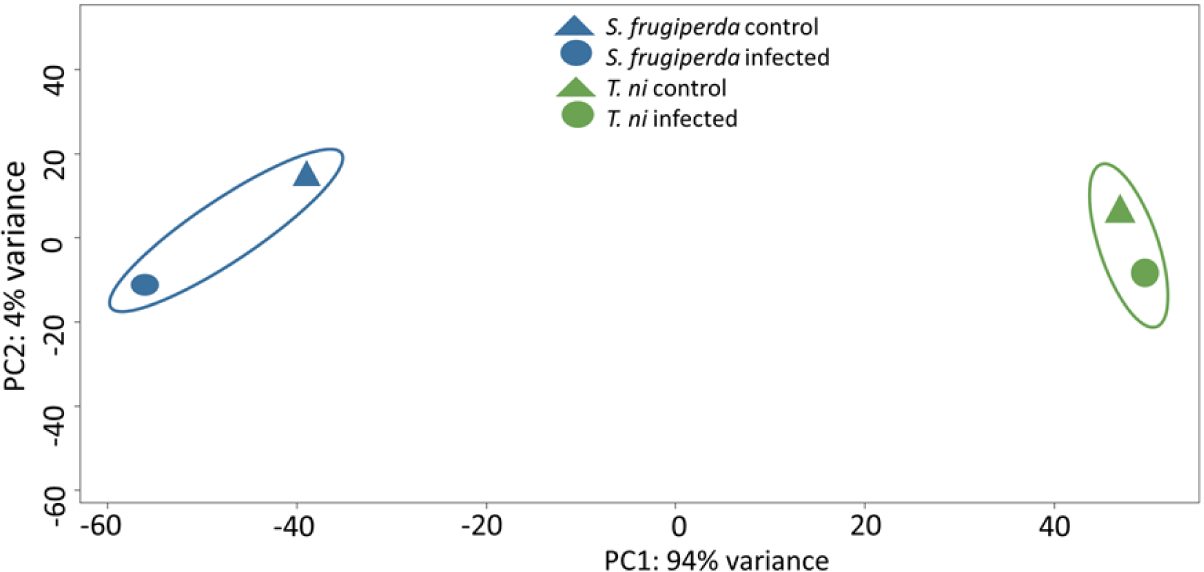
Principle component analysis (PCA) of ortholog gene pairs between *S. frugiperda* and *T. ni*.

**Supplementary Figure 5.**
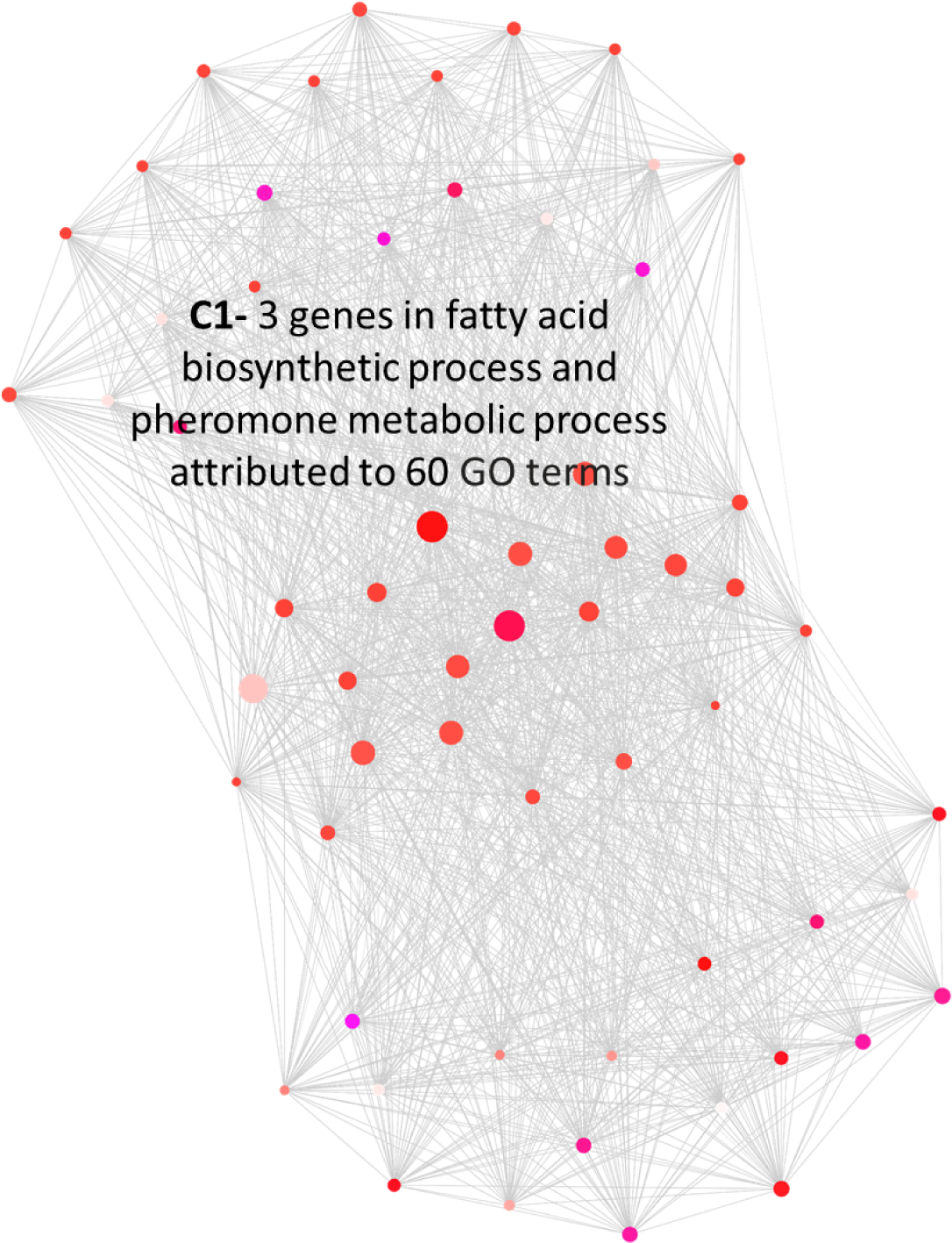
Clustered enriched functional processes among induced *T. ni* transcripts upon AcMNPV infection. The full list of enriched GO terms and GOMCL cluster output are included in the Supplementary Table 5.

**Supplementary Figure 6.**
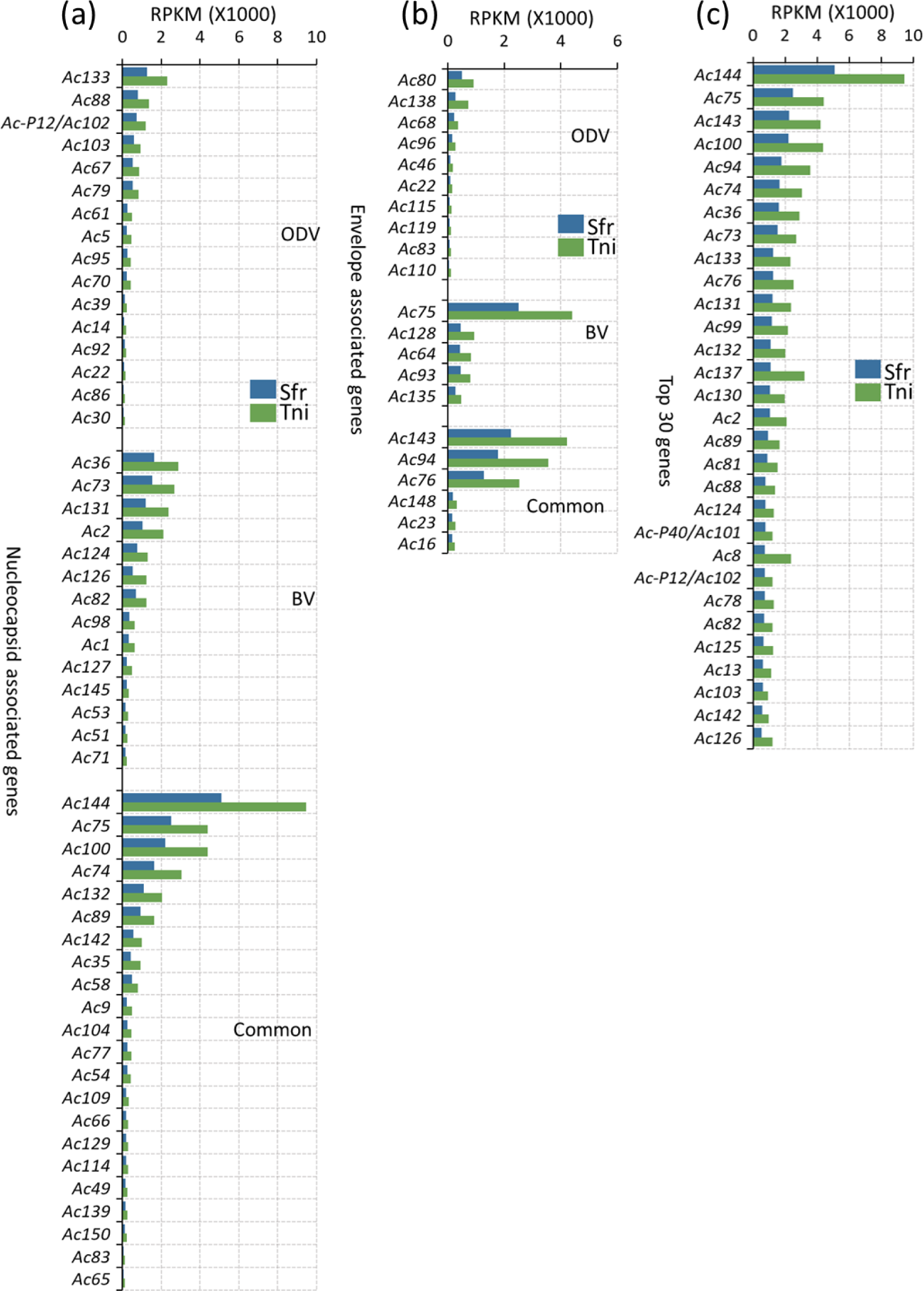
Expression of AcMNPV viral genes in infected *S. frugiperda* and *T. ni*. Genes were classified as nucleocapsid-associated [a] or envelope-associated [b] following Blissard and Theilmenn (2018). Both categories were further divided into genes involved in occlusion derived virus (ODV), budded virus (BV), and common to both the virion types. [c] Top 30 highly abundant viral genes found in *S. frugiperda* and *T. ni* infected larvae.

**Supplementary Table 1.**
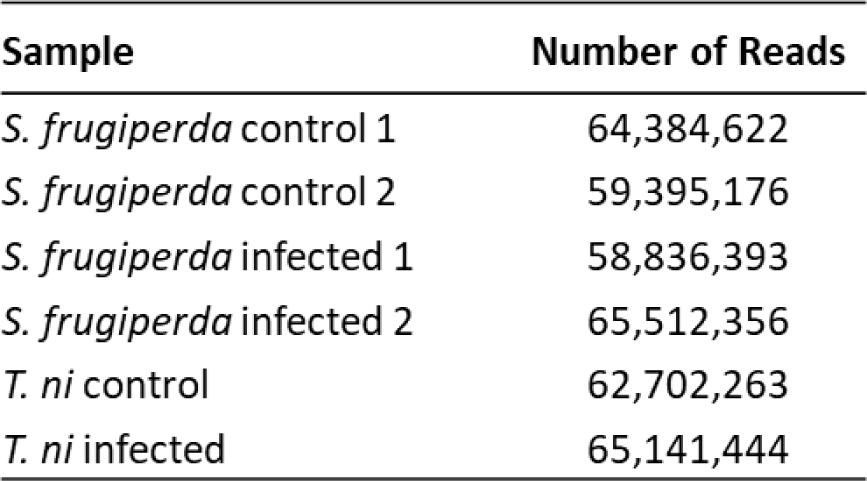
RNAseq data generated for each sample.

| Sample | Number of Reads |
| --- | --- |
| *S. frugiperda* control 1 | 64,384,622 |
| *S. frugiperda* control 2 | 59,395,176 |
| *S. frugiperda* infected 1 | 58,836,393 |
| *S. frugiperda* infected 2 | 65,512,356 |
| *T. ni* control | 62,702,263 |
| *T. ni* infected | 65,141,444 |

**Supplementary Table 2.** [a] Summary of short reads mapped to *S. frugiperda* and *T. ni* reference transcriptomes and [b] percentage of short reads mapped to the AcMNPV viral genome (Maghodia et al., 2014) for control and AcMNPV treated samples.

| (a) | *S. frugiperda* |  | *T. ni* |
| --- | --- | --- | --- |
| Uniquely Mapped (%) | 69.18 |  | 70.70 |
| Reads mapped to multiple contigs (%) | 6.73 |  | 13.28 |
| Total reads mapped (%) | 75.91 |  | 83.98 |
| Total reads processed | 248,128,547 |  | 127,832,614 |

| (b) | *S. frugiperda* |  | *T. ni* |  |
| --- | --- | --- | --- | --- |
|  | control | infected | control | infected |
| Mapped, % | 0.00 | 1.13 | <0.01 | 7.41 |
| Failed to align, % | 100.00 | 98.87 | >99.99 | 92.59 |

**Supplementary Table 3.** Annotation of transcript models with predicted ORFs for *S. frugiperda* and *T. ni* transcriptome assembly.

**Supplementary Table 4.** List of DETs for *S. frugiperda* and *T. ni* in response to AcMNPV infection.

**Supplementary Table 5.** Gene ontology enrichment analysis for DETs for *S. frugiperda* and *T. ni.*

**Supplementary Table 6.** Normalized expression value of AcMNPV genes in *S. frugiperda* and *T. ni* infected hosts.

## References

1. Miller L. The Baculoviruses. Kluwer Academic; 1997.

2. Fuxa JR. Prevalence of Viral Infections in Populations of Fall Armyworm, *Spodoptera frugiperda*, in Southeastern Louisiana. Environ Entomol. 1982;11:239–242.

3. Federici BA. Baculovirus pathogenesis. In: Milleri LK, editor. L.K. Miller (Ed.), The Baculovirus, Plenum, New York. Springer, Boston, MA: Kluwer Academic Publishers; 1997. p. 33–59.

4. Chen Y-R, Zhong S, Fei Z, Hashimoto Y, Xiang JZ, Zhang S, et al. The transcriptome of the baculovirus *Autographa californica* multiple nucleopolyhedrovirus in *Trichoplusia ni* cells. J Virol. 2013;87:6391–405.

5. Blissard GW, Theilmann DA. Baculovirus Entry and Egress from Insect Cells. Annu Rev Virol. 2018;5:113–39.

6. Elderd BD. Developing models of disease transmission: Insights from the ecology of baculovirus-driven systems. PLoS Pathog. 2013;9:e1003372–e1003372.

7. Cory JS, Myers JH. The ecology and evolution of insect baculoviruses. Annu Rev Ecol Evol Syst. 2003;34:239–72.

8. Podgwaite JD, Shields KS, Zerillo RT, Bruen RB. Environmental Persistence of the Nucleopolyhedrosis Virus of the Gypsy Moth, *Lymantria dispar*. Environ Entomol. 1979;8:528–36.

9. Groner A. Specificity and safety of baculoviruses. In: Granados RR, Federici BA, editors. Biology of baculoviruses, Vol. I. Biological Properties and Molecular Biology. CRC Press, Inc.; 1986.

10. Kenoutis C, Efrose RC, Swevers L, Lavdas AA, Gaitanou M, Matsas R, et al. Baculovirus-Mediated Gene Delivery into Mammalian Cells Does Not Alter Their Transcriptional and Differentiating Potential but Is Accompanied by Early Viral Gene Expression. J Virol. 2006;80:4135–46.

11. Condreay J, Kost T. Baculovirus Expression Vectors for Insect and Mammalian Cells. Curr Drug Targets. 2007;8:1126–31.

12. Hu YC. Baculovirus as a highly efficient expression vector in insect and mammalian cells. Acta Pharmacol Sin. 2005;26:405–16.

13. Hu YC. Baculovirus Vectors for Gene Therapy. Adv Virus Res. 2006;68:287–320.

14. Martin DA, SC H, J K, M L-F, RD. P. The compelete DNA sequence of *Autographa californica* nuclear polyhedrosis virus. Virology. 1994;202:586–605.

15. Rohrmann GF. Chapter 12, The AcMNPV genome: Gene content, conservation, and function. Natl Cent Biotechnol Inf. 2013;3rd editio:1–84.

16. Maghodia AB, Jarvis DL, C G. Complete Genome Sequence of the *Autographa californica* Multiple Nucleopolyhedrovirus Strain E2. Genome Announc. 2014;2:6–7.

17. Cory JS, Bishop DHL. Use of Baculoviruses as Biological Insecticides. Mol Biotechnol. 1997;7:303–13.

18. Haase S, Sciocco-Cap A, Romanowski V. Baculovirus Insecticides in Latin America: Historical Overview, Current Status and Future Perspectives. Viruses. 2015;7:2230–67.

19. Belyavskyi M, Braunagel SC, Summers MD. The structural protein ODV-EC27 of *Autographa californica* nucleopolyhedrovirus is a multifunctional viral cyclin. Proc Natl Acad Sci. 1998;95:11205–10.

20. Haas-stapleton EJ, Washburn JO, Volkman LE. Pathogenesis of *Autographa californica* M nucleopolyhedrovirus in fifth instar *Spodoptera frugiperda*. J Gen Virol. 2003;:2033–40.

21. Sutherland DWS, Greene GL. Cultivated and wild host plants. In: P. D. Lingren, & G. U. Greene (Eds.). Suppression and Management of Cabbage Looper Populations. USDA Tech Bull 1684. 1984;:1–13.

22. FAO and CABI. Community-Based Fall Armyworm (*Spodoptera frugiperda*) Monitoring, Early Warning and Management, Training of Trainers Manual, First Edition. 2019.

23. Hardke JT, Leonard BR, Huang F, Jackson RE. Damage and survivorship of fall armyworm (Lepidoptera: Noctuidae) on transgenic field corn expressing *Bacillus thuringiensis* Cry proteins. Crop Prot. 2011;30:168–72.

24. Mota-Sanchez D and JCW. The Arthropod Pesticide Resistance Database. Michigan State Univ. 2020;On-line at:www.pesticideresistance.org.

25. Wang P, Zhao JZ, Rodrigo-Simón A, Kain W, Janmaat AF, Shelton AM, et al. Mechanism of resistance to *Bacillus thuringiensis* toxin Cry1Ac in a greenhouse population of the cabbage looper, *Trichoplusia ni*. Appl Environ Microbiol. 2007;73:1199–207.

26. Lindgren PD, Greene GL. Suppression and Management of Cabbage Looper Populations. 1984.

27. Sparks AN. Review of the biology of the fall armyworm (Lepidoptera, Noctuidae). Florida Entomol. 1979;62:82–87.

28. Richter AR, Fuxa JR, Abdel-Fattah M. Effect of Host Plant on the Susceptibility of *Spodoptera frugiperda* (Lepidoptera: Noctuidae) to a Nuclear Polyhedrosis Virus. Environ Entomol. 1987;16:1004–6.

29. Shorey HH. The Biology of *Trichoplusia ni* (Lepidoptera: Noctuidae). II. Factors Affecting Adult Fecundity and Longevity. Ann Entomol Soc Am. 1963;56:476–480.

30. Pitre HN, Hogg DB. Development of the fall armyworm (Lepidoptera, Noctuidae) on cotton, soybean and corn. J Georg Entomol Soc. 1983;18:182–187.

31. Shorey HH, Andres LA, Hale RL. The Biology of *Trichoplusia ni* (Lepidoptera: Noctuidae). I. Life History and Behavior. Ann Entomol Soc Am. 1962;55:591–597.

32. Anderson R. M., May R. M. The Population Dynamics of Microparasites and Their Invertebrate Hosts. Philos Trans R Soc Lond B Biol Sci. 1981;291:451–524.

33. Salem TZ, Zhang F, Xie Y, Thiem SM. Comprehensive analysis of host gene expression in *Autographa californica* nucleopolyhedrovirus-infected *Spodoptera frugiperda* cells. Virology. 2011;412:167–78.

34. Wei L, Cao L, Miao Y, Wu S, Xu S, Wang R, et al. Transcriptome analysis of *Spodoptera frugiperda* 9 (Sf9) cells infected with baculovirus, AcMNPV or AcMNPV-*Bm*K IT. Biotechnol Lett. 2017;39:1129–39.

35. Chen Y-R, Zhong S, Fei Z, Gao S, Zhang S, Li Z, et al. Transcriptome Responses of the Host *Trichoplusia ni* to Infection by the Baculovirus *Autographa californica* Multiple Nucleopolyhedrovirus. J Virol. 2014;88:13781–97.

36. Shrestha A, Bao K, Chen W, Wang P, Fei Z, Blissard GW. Transcriptional responses of the *Trichoplusia ni* midgut to oral infection by the baculovirus *Autographa californica* Multiple Nucleopolyhedrovirus. J Virol. 2019;93:e00353–19.

37. Xia Q, Zhou Z, Lu C, Cheng D, Dai F, Li B, et al. A draft sequence for the genome of the domesticated silkworm (*Bombyx mori*). Science. 2004;306:1937–40.

38. Pearce SL, Clarke DF, East PD, Elfekih S, Gordon KHJ, Jermiin LS, et al. Genomic innovations, transcriptional plasticity and gene loss underlying the evolution and divergence of two highly polyphagous and invasive *Helicoverpa* pest species. BMC Biol. 2017;15:1–30.

39. Cheng T, Wu J, Wu Y, Chilukuri R V., Huang L, Yamamoto K, et al. Genomic adaptation to polyphagy and insecticides in a major East Asian noctuid pest. Nat Ecol Evol. 2017;1:1747–56.

40. Chen W, Yang X, Tetreau G, Song X, Coutu C, Hegedus D, et al. A high-quality chromosome-level genome assembly of a generalist herbivore, *Trichoplusia ni*. Mol Ecol Resour. 2019;19:485–96.

41. Waterhouse RM, Seppey M, Simao FA, Manni M, Ioannidis P, Klioutchnikov G, et al. BUSCO applications from quality assessments to gene prediction and phylogenomics. Mol Biol Evol. 2018;35:543–8.

42. Mita K, Kasahara M, Sasaki S, Nagayasu Y, Yamada T, Kanamori H, et al. The genome sequence of silkworm, *Bombyx mori*. DNA Res. 2004;11:27–35.

43. Kakumani PK, Malhotra P, Mukherjee SK, Bhatnagar RK. A draft genome assembly of the army worm, *Spodoptera frugiperda*. Genomics. 2014;104:134–43.

44. Legeai F, Gimenez S, Duvic B, Escoubas J-M, Gosselin Grenet A-S, Blanc F, et al. Establishment and analysis of a reference transcriptome for *Spodoptera frugiperda*. BMC Genomics. 2014;15:1–14.

45. Chen Y-R, Zhong S, Fei Z, Hashimoto Y, Xiang JZ, Zhang S, et al. The Transcriptome of the Baculovirus Autographa californica Multiple Nucleopolyhedrovirus in *Trichoplusia ni* Cells. J Virol. 2013;87:6391–405.

46. Chen Y-R, Zhong S, Fei Z, Gao S, Zhang S, Li Z, et al. Transcriptome Responses of the Host *Trichoplusia ni* to Infection by the Baculovirus Autographa californica Multiple Nucleopolyhedrovirus. J Virol. 2014;88:13781–97.

47. Barry MK, Triplett AA, Christensen AC. A peritrophin-like protein expressed in the embryonic tracheae of *Drosophila melanogaster*. Insect Biochem Mol Biol. 1999;29:319–27.

48. Nisole A, Stewart D, Bowman S, Zhang D, Krell PJ, Doucet D, et al. Cloning and characterization of a *Gasp* homolog from the spruce budworm, *Choristoneura fumiferana*, and its putative role in cuticle formation. J Insect Physiol. 2010;56:1427–35.

49. Smith CR, Morandin C, Noureddine M, Pant S. Conserved roles of Osiris genes in insect development, polymorphism and protection. J Evol Biol. 2018;31:516–29.

50. Nagaraj R, Adler PN. Dusky-like functions as a Rab11 effector for the deposition of cuticle during Drosophila bristle development. 2012;139:906–16.

51. Adler PN, Sobala LF, Thom D, Nagaraj R. *dusky-like* is required to maintain the integrity and planar cell polarity of hairs during the development of the Drosophila wing. Dev Biol. 2013;379:76–91.

52. Roch F, Alonso CR, Akam M. *Drosophila miniature* and *dusky* encode ZP proteins required for cytoskeletal reorganisation during wing morphogenesis. J Cell Sci. 2003;116:1199–207.

53. Tonning A, Hemphälä J, Tång E, Nannmark U, Samakovlis C, Uv A. A transient luminal chitinous matrix is required to model epithelial tube diameter in the *Drosophila* trachea. Dev Cell. 2005;9:423–30.

54. Wang S, Jayaram SA, Hemphalä J, Senti K-A, Tsarouhas V, Jin H, et al. Septate-Junction-Dependent Luminal Deposition of Chitin Deacetylases Restricts Tube Elongation in the *Drosophila* Trachea. Curr Biol. 2006;16:180–5.

55. Luschnig S, Ba T, Armbruster K, Krasnow MA. *serpentine* and *vermiform* Encode Matrix Proteins with Chitin Binding and Deacetylation Domains that Limit Tracheal Tube Length in *Drosophila*. Curr Biol. 2006;16:186–94.

56. Petkau G, Wingen C, Jussen LCA, Radtke T, Behr M. Obstructor-A Is Required for Epithelial Extracellular Matrix Dynamics, Exoskeleton Function, and Tubulogenesis. J Biol Chem. 2012;287:21396–405.

57. Zhang L, Iv REW. *uninflatable* encodes a novel ectodermal apical surface protein required for tracheal inflation in *Drosophila*. Dev Biol. 2009;336:201–12.

58. Bunt S, Denholm B, Skaer H. Characterisation of the *Drosophila* procollagen lysyl hydroxylase, *dPlod*. Gene Expr Patterns. 2011;11:72–8.

59. Heikkinen J, Risteli M, Wang C, Latvala J, Rossi M, Valtavaara M, et al. Lysyl Hydroxylase 3 Is a Multifunctional Protein Possessing Collagen Glucosyltransferase Activity *. J Biol Chem. 2000;275:36158–63.

60. Martin GR, Timpl R. Laminin and other basement membrane components. Ann Rev Cell Bioi. 1987;3:57–85.

61. Matsubayashi Y, Louani A, Dragu A, Sánchez-Sánchez BJ, Serna-Morales E, Yolland L, et al. A Moving Source of Matrix Components Is Essential for De Novo Basement Membrane Formation. Curr Biol. 2017;27:3526–34.

62. Chi H-C, Hui C-F. Primary Structure of the Drosophila Laminin B2 Chain and Comparison with Human, Mouse, and Drosophila Laminin B1 and B2 Chains. J Biol Chem. 1989;264:1543–50.

63. Brekken RA, Sage EH, Brekken RA, U EHS. SPARC, a matricellular protein: at the crossroads of cell-matrix communication. Matrix Biol. 2001;19:815–27.

64. Clark CJ, Sage EH. A Prototypic Matricellular Protein in the Tumor Microenvironment— Where There’s SPARC, There’s Fire. J Cell Biochem. 2008;732:721–32.

65. Martinek N, Shahab J, Saathoff M, Ringuette M. Haemocyte-derived SPARC is required for collagen-IV-dependent stability of basal laminae in *Drosophila* embryos. J Cell Sci. 2011;124:1671–80.

66. Shahab J, Baratta C, Scuric B, Godt D, Venken KJT, Ringuette MJ. Loss of SPARC Dysregulates Basal Lamina Assembly to Disrupt Larval Fat Body Homeostasis in *Drosophila melanogaster*. Dev Dyn. 2015;244:540–52.

67. Campbell AG, Fessler LI, Salo T, Fesslerq JH. Papilin: A *Drosophila* Proteoglycan-like Sulfated Glycoprotein from Basement Membranes *. J Biol Chem. 1987;262:17605–12.

68. Hortsch M, Marikar Y, Fishman S, Soneral SN, Dong R, Jacobs JR. The expression of MDP-1, a component of *Drosophila* embryonic basement membranes, is modulated by apoptotic cell death. Int J Dev Biol. 1998;42:33–42.

69. Kramerova IA, Kawaguchi N, Fessler LI, Nelson RE, Chen Y, Kramerov AA, et al. Papilin in development; a pericellular protein with a homology to the ADAMTS metalloproteinases. Development. 2000;127:5475–85.

70. Yamakawa M, Tanaka H. Immune proteins and their gene expression in the silkworm, *Bombyx mori*. Dev Comp Immunol. 1999;23:281–9.

71. Arai I, Ohta M, Suzuki A, Tanaka S, Yoshizawa Y, Sato R. Immunohistochemical Analysis of the Role of Hemocytin in Nodule Formation in the Larvae of the Silkworm, *Bombyx mori*. J Insect Sci. 2013;13:1–13.

72. Ni W, Bao J, Mo B, Liu L, Li T, Pan G, et al. Hemocytin facilitates host immune responses against *Nosema bombycis*. Dev Comp Immunol. 2019;103:103495.

73. Goto A, Kadowaki T, Kitagawa Y. *Drosophila* hemolectin gene is expressed in embryonic and larval hemocytes and its knock down causes bleeding defects. Dev Biol. 2003;264:582–91.

74. Lesch C, Goto A, Lindgren M, Bidla G, Dushay MS, Theopold U. A role for Hemolectin in coagulation and immunity in Drosophila melanogaster. Dev Comp Immunol. 2007;31:1255–63.

75. Kotani E, Yamakawa M, Iwamoto S, Tashiro M, Mori H, Sumida M, et al. Cloning and expression of the gene of hemocytin, an insect humoral lectin which is homologous with the mammalian von Willebrand factor. Biochim Biophys Acta. 1995;1260:245–58.

76. Ponnuvel KM, Yamakawa M. Immune responses against bacterial infection in *Bombyx mori* and regulation of host gene expression. Curr Sci. 2002;83:447–54.

77. Kanost MR, Jiang H. Clip-domain serine proteases as immune factors in insect hemolymph. Curr Opin Insect Sci. 2016;11:47–55.

78. Jagdale S, Bansode S, Joshi R. Insect Proteases: Structural-Functional Outlook in Proteases in physiology and pathology. Springer Nature Singapore; 2017.

79. Houot B, Bousquet F, Ferveur JF. The consequences of regulation of desat1 expression for pheromone emission and detection in *Drosophila melanogaster*. Genetics. 2010;185:1297–309.

80. Moto K, Suzuki MG, Hull JJ, Kurata R, Takahashi S, Yamamoto M, et al. Involvement of a bifunctional fatty-acyl desaturase in the biosynthesis of the silkmoth, *Bombyx mori*, sex pheromone. Proc Natl Acad Sci. 2004;101:8631–6.

81. Dembeck LM, Boroczky K, Huang W, Schal C, Anholt RRH, Mackay TFC. Genetic architecture of natural variation in cuticular hydrocarbon composition in *Drosophila melanogaster*. Elife. 2015;4:1–27.

82. Kefi M, Balabanidou V, Douris V, Lycett G, Feyereisen R, Vontas J. Two functionally distinct CYP4G genes of *Anopheles gambiae* contribute to cuticular hydrocarbon biosynthesis. Insect Biochem Mol Biol. 2019;110:52–9.

83. Qiu Y, Tittiger C, Wicker-Thomas C, Le Goff G, Young S, Wajnberg E, et al. An insect-specific P450 oxidative decarbonylase for cuticular hydrocarbon biosynthesis. Proc Natl Acad Sci. 2012;109:14858–63.

84. Feyereisen R. Origin and evolution of the CYP4G subfamily in insects, cytochrome P450 enzymes involved in cuticular hydrocarbon synthesis. Mol Phylogenet Evol. 2020;143 October 2019:106695.

85. Hayes JD, Flanagan JU, Jowsey IR. Glutathione Transferases. Annu Rev Pharmacol Toxicol. 2005;45:51–88.

86. Da Fonseca RR, Johnson WE, O’Brien SJ, Vasconcelos V, Antunes A. Molecular evolution and the role of oxidative stress in the expansion and functional diversification of cytosolic glutathione transferases. BMC Evol Biol. 2010;10:1–11.

87. Lee M, Yoon CS, Yi J, Cho JR, Kim HS. Cellular immune responses and FAD-glucose dehydrogenase activity of *Mamestra brassicae* (Lepidoptera: Noctuidae) challenged with three species of entomopathogenic fungi. Physiol Entomol. 2005;30:287–92.

88. Pan Q, Shikano I, Felton GW, Liu T-X, Hoover K. Host permissiveness to baculovirus influences time-dependent immune responses and fitness costs. Insect Sci. 2020;00:1–12.

89. Katsuma S, Kawaoka S, Mita K, Shimada T. Genome-wide survey for baculoviral host homologs using the *Bombyx* genome sequence. Insect Biochem Mol Biol. 2008;38:1080–6.

90. Gilbert C, Chateigner A, Ernenwein L, Barbe V, Annie B, Herniou2 EA, et al. Population genomics supports baculoviruses as vectors of horizontal transfer of insect transposons. Nat Commun. 2014;5:1–9.

91. Clem RJ, Passarelli AL. Baculoviruses: Sophisticated Pathogens of Insects. PLoS Pathog. 2013;9:11–4.

92. Choi JY, Roh JY, Wang Y, Zhen Z, Tao XY, Lee JH, et al. Analysis of genes expression of *Spodoptera exigua* Larvae upon AcMNPV infection. PLoS One. 2012;7:1–10.

93. Guarino LA, Dong WEN, Xu BIN, Broussard DR, Davis RW, Jarvis DL. Baculovirus Phosphoprotein pp3l Is Associated with Virogenic Stroma. J Virol. 1992;66:7113–20.

94. Guo ZJ, Qiu LH, An SH, Yao Q, Park EY, Chen KP, et al. Open reading frame 60 of the *Bombyx mori* nucleopolyhedrovirus plays a role in budded virus production. Virus Res. 2010;151:185–91.

95. Wei D, Wang Y, Zhang X, Hu Z, Yuan M, Yang K. *Autographa californica Nucleopolyhedrovirus* Ac76: a Dimeric Type II Integral Membrane Protein That Contains an Inner Nuclear Membrane-Sorting Motif. J Virol. 2014;88:1090–103.

96. Hu Z, Yuan M, Wu W, Liu C, Yang K, Pang Y. *Autographa californica* Multiple Nucleopolyhedrovirus *ac76* Is Involved in Intranuclear Microvesicle Formation. J Virol. 2010;84:7437–47.

97. Chen L, Yang R, Hu X, Xiang X, Yu S, Wu X. The formation of occlusion-derived virus is affected by the expression level of ODV-E25. Virus Res. 2013;173:404–14.

98. Chen L, Hu X, Xiang X, Yu S, Yang R, Wu X. *Autographa californica* multiple nucleopolyhedrovirus *odv-e25* (Ac94) is required for budded virus infectivity and occlusion-derived virus formation. Arch Virol. 2012;157:617–25.

99. Wang M, Tuladhar E, Shen S, Wang H, van Oers MM, Vlak JM, et al. Specificity of Baculovirus P6.9 Basic DNA-Binding Proteins and Critical Role of the C Terminus in Virion Formation. J Virol. 2010;84:8821–8.

100. McCarthy CB, Theilmann DA. AcMNPV *ac143* (*odv-e18*) is essential for mediating budded virus production and is the 30th baculovirus core gene. Virology. 2008;375:277–91.

101. Braunagel SC, He H, Ramamurthy P, Summers MD. Transcription, Translation, and Cellular Localization of Three *Autographa californica* Nuclear Polyhedrosis Virus Structural Proteins: ODV-E18, ODV-E35, and ODV-EC27. Virology. 1996;114:100–14.

102. Gross CH, Russell RLQ, Rohrmann GF. *Orgyia pseudotsugata* baculovirus p10 and polyhedron envelope protein genes: Analysis of their relative expression levels and role in polyhedron structure. J Gen Virol. 1994;75:1115–23.

103. McCarthy CB, Dai X, Donly C, Theilmann DA. *Autographa californica* multiple nucleopolyhedrovirus *ac142*, a core gene that is essential for BV production and ODV envelopment. Virology. 2008;372:325–39.

104. Vanarsdall AL, Pearson MN, Rohrmann GF. Characterization of baculovirus constructs lacking either the Ac 101, Ac 142, or the Ac 144 open reading frame. Virology. 2007;367:187–95.

105. Hawtin RE, Zarkowska T, Arnold K, Thomas CJ, Gooday GW, King LA, et al. Liquefaction of *Autographa californica* nucleopolyhedrovirus-infected insects is dependent on the integrity of virus-encoded chitinase and cathepsin genes. Virology. 1997;238:243–53.

106. Slack JM, Kuzio J, Faulkner P. Characterization of *v-cath*, a cathepsin L-like proteinase expressed by the baculovirus *Autographa californica* multiple nuclear polyhedrosis virus. J Gen Virol. 1995;76:1091–8.

107. Ishimwe E, Hodgson JJ, Clem RJ, Passarelli AL. Reaching the melting point: Degradative enzymes and protease inhibitors involved in baculovirus infection and dissemination. Virology. 2015;479–480:637–49.

108. Lyupina Y V, Dmitrieva SB, Timokhova A V, Beljelarskaya SN, Zatsepina OG, Evgen MB, et al. An important role of the heat shock response in infected cells for replication of baculoviruses. Virology. 2010;406:336–41.

109. Mu J, Zhang Y, Hu Y, Hu X, Zhou Y, Zhao H, et al. Autographa californica Multiple Nucleopolyhedrovirus Ac34 Protein Retains Cellular Actin-Related Protein 2/3 Complex in the Nucleus by Subversion of CRM1-Dependent Nuclear Export. PLoS Pathog. 2016;12:1–22.

110. Lyupina YV, Orlova OV, Abaturova SB, Beljelarskaya SN, Lavrov AN, Mikhailov VS. Egress of budded virions of *Autographa californica* nucleopolyhedrovirus does not require activity of *Spodoptera frugiperda* HSP/HSC70 chaperones. Virus Res. 2014;192:1–5.

111. Heaton NS, Randall G. Multifaceted roles for lipids in viral infection. Trends Microbiol. 2011;19:368–75.

112. Ohkawa T, Volkman LE, Welch MD. Actin-based motility drives baculovirus transit to the nucleus and cell surface. J Cell Biol. 2010;190:187–95.

113. Qiu J, Tang Z, Yuan M, Wu W, Yang K. The 91-205 amino acid region of AcMNPV ORF34 (Ac34), which comprises a potential C3H zinc finger, is required for its nuclear localization and optimal virus multiplication. Virus Res. 2017;228:79–89.

114. Cai Y, Long Z, Qiu J, Yuan M, Li G, Yang K. An *ac34* Deletion Mutant of *Autographa californica* Nucleopolyhedrovirus Exhibits Delayed Late Gene Expression and a Lack of Virulence *In Vivo*. J Virol. 2012;86:10432–43.

115. Kelly KK, Meadows SM, Cripps RM. Drosophila MEF2 is a direct regulator of Actin57B transcription in cardiac, skeletal, and visceral muscle lineages. Mech Dev. 2002;110:39–50.

116. Vierstraete E, Cerstiaens A, Baggerman G, Bergh G Van Den, Loof A De, Schoofs L. Proteomics in *Drosophila melanogaster*: first 2D database of larval hemolymph proteins. Biochem Biophys Res Commun. 2003;304:831–8.

117. Cossart P. Actin-based motility of pathogens: The Arp2/3 complex is a central player. Cell Microbiol. 2000;2:195–205.

118. Devitt A, Marshall LJ. The innate immune system and the clearance of apoptotic cells. J Leukoc Biol. 2011;90:447–57.

119. Krieser RJ, Eastman A. Cleavage and nuclear translocation of the caspase 3 substrate Rho GDP-dissociation inhibitor, D4-GDI, during apoptosis. Cell Death Differ. 1999;6:412–9.

120. Inbal B, Bialik S, Sabanay I, Shani G, Kimchi A. DAP kinase and DRP-1 mediate membrane blebbing and the formation of autophagic vesicles during programmed cell death. J Cell Biol. 2002;157:455–68.

121. Noriegaa FG, Ribeirob JMC, Koenerc JF, Valenzuelab JG, Hernandez-Martinez S, Pham VM, et al. Comparative genomics of insect juvenile hormone biosynthesis. Insect Biochem Mol Biol. 2006;36:366–74.

122. Bailey D, Basar MA, Nag S, Bondhu N, Teng S, Duttaroy A. The essential requirement of an animal heme peroxidase protein during the wing maturation process in Drosophila. BMC Dev Biol. 2017;:1–11.

123. Reilly DRO, Miller LK. A Baculovirus Blocks Insect Molting by Producing. 1989;245:1110–2.

124. Rao R, Fiandra L, Giordana B, De Eguileor M, Congiu T, Burlini N, et al. AcMNPV ChiA protein disrupts the peritrophic membrane and alters midgut physiology of *Bombyx mori* larvae. Insect Biochem Mol Biol. 2004;34:1205–13.

125. Erlandson MA, Toprak U, Hegedus DD. Role of the peritrophic matrix in insect-pathogen interactions. J Insect Physiol. 2019;117 June:103894.

126. He L, Li N, Chen Y, Liu S. Regulation of Chitinase in *Spodoptera exigua* (Hübner) (Lepidoptera:Noctuidae) During Infection by *Heliothis virescens*. Front Physiol. 2020;11:1–11.

127. Su C, Tu G, Huang S, Yang Q. Genome-wide analysis of chitinase genes and their varied functions in larval moult, pupation and eclosion in the rice striped stem borer, *Chilo suppressalis*. Insect Mol Biol. 2016;25:401–12.

128. Caragata EP, Pais FS, Baton LA, Silva JBL, Sorgine MHF, Moreira LA. The transcriptome of the mosquito *Aedes fluviatilis* (Diptera: Culicidae), and transcriptional changes associated with its native *Wolbachia* infection. BMC Genomics. 2017;18:1–19.

129. Zhu Q, Arakane Y, Banerjee D, Beeman RW, Kramer KJ, Muthukrishnan S. Domain organization and phylogenetic analysis of the chitinase-like family of proteins in three species of insects. Insect Biochem Mol Biol. 2008;38:452–66.

130. Hawtin RE, Arnold K, Ayres MD, Zanotto PMD, Howard SC, Gooday GW, et al. Identification and Preliminary Characterization of a Chitinase Gene in the *Autographa californica* Nuclear Polyhedrosis-Virus Genome. Virology. 1995;212:673–85.

131. Daimon T, Katsuma S, Kang WK, Shimada T. Comparative studies of *Bombyx mori* nucleopolyhedrovirus *chitinase* and its host ortholog, *BmChi-h*. Biochem Biophys Res Commun. 2006;345:825–33.

132. Beck K, Hunter I, Engel J. Structure and function of laminin: anatomy of a multidomain glycoprotein. FASEB J. 1990;4:148–60.

133. Keeley DP, Hastie E, Jayadev R, Kelley LC, Chi Q, Payne SG, et al. Comprehensive Endogenous Tagging of Basement Membrane Components Reveals Dynamic Movement within the Matrix Scaffolding. Dev Cell. 2020;54:60–74.e7.

134. Means JC, Passarelli AL. Viral fibroblast growth factor, matrix metalloproteases, and caspases are associated with enhancing systemic infection by baculoviruses. Proc Natl Acad Sci. 2010;107:9825–30.

135. Monsma SA, Oomens AGP, Blissard GW. The GP64 Envelope Fusion Protein Is an Essential Baculovirus Protein Required for Cell-to-Cell Transmission of Infection. J Virol. 1996;70:4607–16.

136. Passarelli AL. Barriers to success: How baculoviruses establish efficient systemic infections. Virology. 2012;411:383–92.

137. Li J, Sun Y, Li Y, Liu X, Yue Q, Li Z. Inhibition of cellular fatty acid synthase impairs replication of budded virions of Autographa californica multiple nucleopolyhedrovirus in *Spodoptera frugiperda* cells. Virus Res. 2018;252 March:41–7.

138. Munger J, Bennett BD, Parikh A, Feng X, Rabitz HA, Shenk T, et al. Systems-level metabolic flux profiling identifies fatty acid synthesis as a target for antiviral therapy. Nat Biotechnol. 2010;26:1179–86.

139. Breitenbach JE, Shelby KS, Popham HJR. Baculovirus induced transcripts in hemocytes from the larvae of *Heliothis virescens*. Viruses. 2011;3:2047–64.

140. Bousquet F, Ferveur JF. desat1 A Swiss army knife for pheromonal communication and reproduction? Fly (Austin). 2012;6:102–7.

141. Köhler K, Brunner E, Xue LG, Boucke K, Greber UF, Mohanty S, et al. A combined proteomic and genetic analysis identifies a role for the lipid desaturase Desat1 in starvation-induced autophagy in Drosophila. Autophagy. 2009;5:980–90.

142. He C, Klionsky DJ. Regulation Mechanisms and Signaling Pathways of Autophagy. Annu Rev Genet. 2009;43:67–93.

143. Roxstrom-Lindquist K, Terenius O, Faye I. Parasite-specific immune response in adult *Drosophila melanogaster*: a genomic study. EMBO Rep. 2004;5:207–12.

144. Cerenius L, Kawabata S ichiro, Lee BL, Nonaka M, Söderhäll K. Proteolytic cascades and their involvement in invertebrate immunity. Trends Biochem Sci. 2010;35:575–83.

145. Lavine MD, Strand MR. Insect Hemocytes and Their Role in Immunity. Insect Immunol. 2008;32:25–47.

146. Zhao P, Wang GH, Dong ZM, Duan J, Xu PZ, Cheng TC, et al. Genome-wide identification and expression analysis of serine proteases and homologs in the silkworm *Bombyx mori*. BMC Genomics. 2010;11:1–11.

147. Tang H, Kambris Z, Lemaitre B, Hashimoto C. Two Proteases Defining a Melanization Cascade in the Immune System of Drosophila*. J Biol Chem. 2006;281:28097–104.

148. Washburn JO, Haas-Stapleton EJ, Tan FF, Beckage NE, Volkman LE. Co-infection of *Manduca sexta* larvae with polydnavirus from *Cotesia congregata* increases susceptibility to fatal infection by *Autographa californica* M Nucleopolyhedrovirus. J Insect Physiol. 2000;46:179–90.

149. Engelhard EK, Kam-Morgan LN, Washburn JO, Volkman LE. The insect tracheal system: a conduit for the systemic spread of *Autographa californica* M nuclear polyhedrosis virus. Proc Natl Acad Sci. 1994;91:3224–7.

150. Monteiro F, Carinhas N, Carrondo MJT, Bernal V. Toward system-level understanding of baculovirus–host cell interactions: from molecular fundamental studies to large-scale proteomics approaches. Front Microbiol. 2012;3:1–16.

151. Lavington E, Cogni R, Kuczynski C, Koury S, Behrman EL, O’brien KR, et al. A small system-high-resolution study of metabolic adaptation in the central metabolic pathway to temperate climates in *Drosophila melanogaster*. Mol Biol Evol. 2014;31:2032–41.

152. Wise EM, Ball EG. Malic Enzyme and Lipogenesis. Proc Natl Acad Sci. 1964;52:1255–63.

153. Merritt TJS, Duvernell D, Eanes WF. Natural and synthetic alleles provide complementary insights into the nature of selection acting on the *Men* polymorphism of *Drosophila melanogaster*. Genetics. 2005;171:1707–18.

154. Geer BW, Krochko D, Williamson JH. Ontogeny, cell distribution, and the physiological role of NADP-malic enzyme in *Drosophila melanogaster*. Biochem Genet. 1979;17:867–79.

155. Schwarz KB. Oxidative stress during viral infection: a review. Free Radic Biol Med. 1996;21:641–9.

156. Camini FC, Lopes C, Magalha DB. Implications of oxidative stress on viral pathogenesis. Arch Virol. 2017;162:907–17.

157. Wertheim B, van Baalen E-JA, Dicke M, Vet LEM. Pheromone-mediated aggregation in nonsocial arthropods: An Evolutionary Ecological Perspective. Annu Rev Entomol. 2005;50:321–46.

158. Trienens M, Rohlfs M. A Potential Collective Defense of Drosophila Larvae Against the Invasion of a Harmful Fungus. Front Ecol Evol. 2020;8:1–10.

159. Gomez-Diaz C, Bargeton B, Abuin L, Bukar N, Reina JH, Bartoi T, et al. A CD36 ectodomain mediates insect pheromone detection via a putative tunnelling mechanism. Nat Commun. 2016;7:1–17.

160. Benton R, Vannice KS, Vosshall LB. An essential role for a CD36-related receptor in pheromone detection in *Drosophila*. Nature. 2007;450:289–93.

161. Mast JD, De Moraes CM, Alborn HT, Lavis LD, Stern DL. Evolved differences in larval social behavior mediated by novel pheromones. Elife. 2014;3:e04205.

162. Sabatier L, Jouanguy E, Dostert C, Zachary D, Dimarcq JL, Bulet P, et al. Pherokine-2 and -3: Two *Drosophila* molecules related to pheromone/odor-binding proteins induced by viral and bacterial infections. Eur J Biochem. 2003;270:3398–407.

163. Labeur C, Dallerac R, Wicker-Thomas C. Involvement of *desat1* gene in the control of *Drosophila melanogaster* pheromone biosynthesis. Genetica. 2002;114:269–74.

164. Ueyama M, Chertemps T, Labeur C, Wicker-Thomas C. Mutations in the *desat1* gene reduces the production of courtship stimulatory pheromones through a marked effect on fatty acids in *Drosophila melanogaster*. Insect Biochem Mol Biol. 2005;35:911–20.

165. Bousquet F, Nojima T, Houot B, Chauvel I, Chaudy S, Dupas S, et al. Expression of a desaturase gene, *desat1*, in neural and nonneural tissues separately affects perception and emission of sex pheromones in *Drosophila*. Proc Natl Acad Sci. 2012;109:249–54.

166. Sim S, Ramirez L, Dimopoulos G. Dengue Virus Infection of the *Aedes aegypti* Salivary Gland and Chemosensory Apparatus Induces Genes that Modulate Infection and Blood-Feeding Behavior. PLoS Pathog. 2012;8:e1002631.

167. Bartholomay LC, Cho WL, Rocheleau TA, Boyle JP, Beck ET, Fuchs JF, et al. Description of the transcriptomes of immune response-activated hemocytes from the mosquito vectors *Aedes aegypti* and *Armigeres subalbatus*. Infect Immun. 2004;72:4114–26.

168. Wang Y, Oberley LW, Murhammer DW. Evidence of oxidative stress following the viral infection of two lepidopteran insect cell lines. Free Radic Biol Med. 2001;31:1448–55.

169. Schieber M, Chandel NS. ROS function in redox signaling. Curr Biol. 2014;24:453–62.

170. Shi A, Hu Z, Zuo Y, Wang Y, Wu W, Yuan M, et al. Autographa californica Multiple Nucleopolyhedrovirus *ac75* Is Required for the Nuclear Egress of Nucleocapsids and Intranuclear Microvesicle Formation. J Virol. 2018;92:e01509–17.

171. Guo Y, Fu S, Li L. Autographa californica multiple nucleopolyhedrovirus ac75 is required for egress of nucleocapsids from the nucleus and formation of de novo intranuclear membrane microvesicles. PLoS One. 2017;12:e0185630.

172. Kadlec J, Loureiro S, Abrescia NGA, Stuart DI, Jones IM. The postfusion structure of baculovirus gp64 supports a unified view of viral fusion machines. Nat Struct Mol Biol. 2008;15:1024–30.

173. Dong S, Blissard GW. Functional Analysis of the *Autographa californica* Multiple Nucleopolyhedrovirus GP64 Terminal Fusion Loops and Interactions with Membranes. J Virol. 2012;86:9617–28.

174. Sun X, Belouzard S, Whittaker GR. Molecular Architecture of the Bipartite Fusion Loops of Vesicular Stomatitis Virus Glycoprotein G, a Class III Viral Fusion Protein. J Biol Chem. 2008;283:6418–27.

175. Wang M, Wang J, Yin F, Tan Y, Deng F, Chen X, et al. Unraveling the Entry Mechanism of Baculoviruses and Its Evolutionary Implications. J Virol. 2014;88:2301–11.

176. Pearson MN, Rohrmann GF. Transfer, Incorporation, and Substitution of Envelope Fusion Proteins among Members of the *Baculoviridae*, *Orthomyxoviridae*, and *Metaviridae* (Insect Retrovirus) Families. J Virol. 2002;76:5301–4.

177. Braunagel SC, Parr R, Belyavskyi M, Summers MD. *Autographa californica* nucleopolyhedrovirus infection results in Sf9 cell cycle arrest at G2/M phase. Virology. 1998;244:195–211.

178. Ishidate T, Elewa A, Kim S, Mello CC, Shirayama M. Divide and differentiate CDK / Cyclins and the art of development. Cell Cycle. 2014;13:1384–91.

179. O’Reilly DR, Miller LK. Regulation of expression of a baculovirus ecdysteroid UDPglucosyltransferase gene. J Virol. 1990;64:1321–8.

180. Hoover K, Grove M, Gardner M, Hughes DP, Mcneil J, Slavicek J. A Gene for an Extended Phenotype. Science. 2011;333:1401.

181. Ros VID, Van Houte S, Hemerik L, Van Oers MM. Baculovirus-induced tree-top disease: How extended is the role of *egt* as a gene for the extended phenotype? Mol Ecol. 2015;24:249–58.

182. Van Houte S, Ros VID, Van Oers MM. Hyperactivity and tree-top disease induced by the baculovirus AcMNPV in *Spodoptera exigua* larvae are governed by independent mechanisms. Naturwissenschaften. 2014;101:347–50.

183. Herrero S, Ansems M, Van Oers MM, Vlak JM, Bakker PL, de Maagd RA. REPAT, a new family of proteins induced by bacterial toxins and baculovirus infection in *Spodoptera exigua*. Insect Biochem Mol Biol. 2007;37:1109–18.

184. Merzendorfer H. Chitin synthesis inhibitors: Old molecules and new developments. Insect Sci. 2013;20:121–38.

185. Zhu KY, Merzendorfer H, Zhang W, Zhang J, Muthukrishnan S. Biosynthesis, Turnover, and Functions of Chitin in Insects. Annu Rev Entomol. 2016;61:177–96.

186. Douris V, Steinbach D, Panteleri R, Livadaras I, Pickett JA, Van Leeuwen T, et al. Resistance mutation conserved between insects and mites unravels the benzoylurea insecticide mode of action on chitin biosynthesis. Proc Natl Acad Sci. 2016;113:14692–7.

187. Moussian B, Schwarz H, Bartoszewski S, Nu C. Involvement of Chitin in Exoskeleton Morphogenesis in *Drosophila melanogaster*. J Morphol. 2005;264:117–30.

188. Cook DR, Leonard BR, Gore J. Field and Laboratory Performance of Novel Insecticides Against Armyworms (Lepidoptera: Noctuidae). Florida Entomol. 2006;87:433–9.

189. IRAC South Africa. Integrated Pest Management (IPM) & Insect Resistance Management (IRM) for Fall Armyworm in South African Maize. 2018;:1–21.

190. Powell GF, Ward DA, Prescott MC, Spiller DG, White MRH, Turner PC, et al. The molecular action of the novel insecticide, Pyridalyl. Insect Biochem Mol Biol. 2011;41:459–69.

191. San Miguel K, Scott JG. The next generation of insecticides: DsRNA is stable as a foliar-applied insecticide. Pest Manag Sci. 2016;72:801–9.

192. Wilson K, Grzywacz D, Curcic I, Scoates F, Harper K, Rice A, et al. A novel formulation technology for baculoviruses protects biopesticide from degradation by ultraviolet radiation. Sci Rep. 2020;10:13301.

193. Shim HJ, Choi JY, Wang Y, Tao XY, Liu Q, Roh JY, et al. Neurobactrus, a novel, highly effective, and environmentally friendly recombinant baculovirus insecticide. Appl Environ Microbiol. 2013;79:141–9.

194. Cory JS, Hirst ML, Williams T, Hails RS, Goulson D, Green BM, et al. Field trial of a genetically improved baculovirus insecticide. Nature. 1994;370:138–40.

195. Bentivenha JPF, Rodrigues JG, Lima MF, Marçon P, Popham HJR, Omoto C. Baseline Susceptibility of *Spodoptera frugiperda* (Lepidoptera: Noctuidae) to SfMNPV and Evaluation of Cross-Resistance to Major Insecticides and Bt Proteins. J Econ Entomol. 2019;112:91–8.

196. Das S, Goswami A, Debnath N. Application of baculoviruses as biopesticides and the possibilities of nanoparticle mediated delivery in Nano-Biopesticides Today and Future Perspectives. Elsevier Inc.; 2019.

197. Liu Z, Wang X, Dai Y, Wei X, Ni M, Zhang L, et al. Expressing double-stranded RNAs of insect hormone-related genes enhances baculovirus insecticidal activity. Int J Mol Sci. 2019;20:1–13.

198. Bonning BC, Hirst M, Possee RD, Hammock BD. Further development of a recombinant baculovirus insecticide expressing the enzyme juvenile hormone esterase from *Heliothis virescens*. Insect Biochem Mol Biol. 1992;22:453–8.

199. Chen E, Kolosov D, O’Donnell MJ, Erlandson MA, McNeil JN, Donly C. The effect of diet on midgut and resulting changes in infectiousness of AcMNPV baculovirus in the cabbage looper, *Trichoplusia ni*. Front Physiol. 2018;9 OCT:1–11.

200. Collett D. Modelling binary data. Boca Raton, FL: Chapman & Hall/CRC; 2003.

201. Team R core. R: A Language and Environment for Statistical Computing. 2018.

202. Plummer M. JAGS: A Program for Analysis of Bayesian Graphical Models Using Gibbs Sampling. Proc DSC. 2003;:1–10.

203. Su Y, Yajima M. Package ‘R2jags.’ 2015.

204. Link WA, Eaton MJ. On thinning of chains in MCMC. Methods Ecol Evol. 2012;3:112–115.

205. Gelman A, Carlin JB, Stern HS, Rubin DB. Bayesian data analysis. Taylor & Francis; 2014.

206. Gelman A, Meng XL, Stern H. Posterior predictive assessment of model fitness via realized discrepancies. Stat Sin. 1996;6:733–760.

207. Kéry M. Introduction to WinBUGS for ecologists. 2010.

208. Fan L, Wang G, Hu W, Pantha P, Tran K-N, Zhang H, et al. Transcriptomic view of survival during early seedling growth of the extremophyte *Haloxylon ammodendron*. Plant Physiol Biochem. 2018;132:475–89.

209. Grabherr MG, Haas BJ, Yassour M, Levin JZ, Thompson DA, Amit I, et al. Full-length transcriptome assembly from RNA-Seq data without a reference genome. Nat Biotechnol. 2011;29:644–52.

210. Oh D, Barkla BJ, Vera-estrella R, Pantoja O, Lee S, Bohnert HJ, et al. Cell type-specific responses to salinity–the epidermal bladder cell transcriptome of *Mesembryanthemum crystallinum*. New Phytol. 2015;207:627–44.

211. Li W, Godzik A. Cd-hit: A fast program for clustering and comparing large sets of protein or nucleotide sequences. Bioinformatics. 2006;22:1658–9.

212. Simão FA, Waterhouse RM, Ioannidis P, Kriventseva E V., Zdobnov EM. BUSCO: Assessing genome assembly and annotation completeness with single-copy orthologs. Bioinformatics. 2015;31:3210–2.

213. Langmead B, Trapnell C, Pop M, Salzberg SL. Ultrafast and memory-efficient alignment of short DNA sequences to the human genome. Genome Biol. 2009;10:R25.

214. Tarazona S, Furió-Tarí P, Turrà D, Pietro A Di, Nueda MJ, Ferrer A, et al. Data quality aware analysis of differential expression in RNA-seq with NOISeq R/Bioc package. Nucleic Acids Res. 2015;43:e140.

215. Maere S, Heymans K, Kuiper M. BiNGO: A Cytoscape plugin to assess overrepresentation of Gene Ontology categories in Biological Networks. Bioinformatics. 2005;21:3448–9.

216. Wang G, Oh D, Dassanayake M. GOMCL: a toolkit to cluster, evaluate, and extract non-redundant associations of Gene Ontology-based functions. BMC Bioinformatics. 2020;21:1–9.

217. Shrestha A, Bao K, Chen Y-R, Chen W, Wang P, Fei Z, et al. Global Analysis of Baculovirus *Autographa californica* Multiple Nucleopolyhedrovirus Gene Expression in the Midgut of the Lepidopteran Host *Trichoplusia ni*. J Virol. 2018;92:e01277–18.

